# Identifying adaptive variation in spatially structured populations using low-coverage whole-genome sequencing data

**DOI:** 10.64898/2026.01.19.700474

**Authors:** Nikunj Goel, Christen M. Bossu, Seorim Yi, Timothy M. Brown, Erica C. N. Robertson, Peri E. Bolton, Ben J. Vernasco, Reza Goljani Amirkhiz, Erika Zavaleta, Kristen C. Ruegg, Mevin B. Hooten

## Abstract

Successful implementation of evolutionary programs to rescue climatically threatened species requires identification of adaptive variation. Although many genotype-environment association methods have been successful in identifying adaptive variation, current approaches can be improved in two important aspects. First, most existing methods do not account for genotype uncertainty in widely available low-coverage whole-genome sequencing data. Researchers often restrict analysis to loci for which genotypes can be inferred reliably or call the most probable genotype, allowing the use of genotype-based methods. However, discarding data and false genotype calls increase the uncertainty in estimates of genetic variation and can introduce systematic biases. Second, most methods use phenomenological approaches, such as logistic regression, to partition estimated variation into adaptive and non-adaptive components. Consequently, current approaches may fail to account for evolutionary processes, such as migration-selection balance. Structured migration between climatically disparate locations can produce deviations from a smooth S-shape response curve, which can be difficult to accommodate using generalized linear models. To overcome these challenges, we developed a method that accounts for genotype uncertainty in sequencing data and propagates this uncertainty to inform the parameters of an evolutionary model. A key feature of this model is that it describes mechanistically how genetic variation arises from joint interactions between local adaptation, structured migration, mutation, and drift. Our synthetic simulation tests reveal that accounting for genotype uncertainty and structured migration substantially reduces false negatives. We also applied our approach to analyze data on North American rosy-finches (3.7 million SNPs), a high-alpine, climatically threatened clade of bird species.

## Introduction

Contemporary climate change is expected to drive many species to extinction, especially those with limited dispersal capacity to track their environmental niches (Schloss et al. 2012) and those that occupy shrinking suitable habitats, such as sky islands (Freeman et al. 2018). Conservationists argue that evolutionary management policies, such as assisted gene flow and genome-editing (Kosch et al. 2022; Phelps et al. 2020), may provide a viable path to rescue these species that are otherwise likely to go extinct (Harrisson et al. 2014; Iverson 2024). A key prerequisite for the successful implementation of these management programs is identifying adaptive loci, which may enable managers to target specific regions of the genome for amelioration.

Over the past two decades, several statistical methods have been developed to identify adaptive genomic regions in climatically threatened species (see Rellstab et al. 2015 for a review of studies). Broadly, these statistical methods, referred to as genotype-environment association methods, work in three steps (Goel et al. 2025). First, individuals of a species are sampled across its geographical range and are subsequently sequenced in the lab to estimate standing genetic variation. Second, the estimated genetic variation is partitioned into adaptive and non-adaptive (neutral) components. The adaptive variation corresponds to clinal patterns in allele frequencies or genotypes that are congruent with environmental variation. The non-adaptive variation corresponds to genetic patterns that arise from neutral evolutionary processes, such as mutation, migration, and genetic drift. Finally, a statistical technique is used to evaluate the relative contribution of adaptive and non-adaptive evolutionary forces in determining variation at a given locus. Despite substantial methodological progress in ecological genomics, the capabilities of current genotype-environment association methods can be further expanded to enable more robust statistical inferences.

When estimating genetic variation using low-coverage whole-genome sequencing data, many genotype-environment association methods do not faithfully account for, and propagate, genotype uncertainty. Most current methods that use individual-level sequencing data require genotypes as inputs to estimate genetic variation (De Villemereuil and Gaggiotti 2015; Frichot et al. 2013; Günther and Coop 2013). However, in low-coverage sequencing data, an individual’s genotype can only be inferred probabilistically (Nielsen et al. 2012; Nielsen et al. 2011). To address this issue, current genotype-environment association methods take the following routes. Researchers either use the most probable genotype, impute genotypes based on linkage maps (Li et al. 2011; Lou et al. 2021), or analyze only a subset of loci for which genotypes can be called with high certainty (Andrews et al. 2023). However, this approach is suboptimal because it can introduce systematic biases from false genotype calls and result in loss of information due to reduced sample size and limited genomic coverage (Han et al. 2014). Alternatively, some researchers first estimate allele frequencies that account for sampling and genotype uncertainty using statistical software, such as ANGSD (Korneliussen et al. 2014), and then use point estimates (e.g., the mean) as inputs in the downstream analysis (Capblancq et al. 2018; Forester et al. 2018). However, this approach is also suboptimal because statistical models underestimate uncertainty in relevant downstream parameters (Fumagalli et al. 2013). An improved approach to accommodate low-coverage whole-genome sequencing data is to simultaneously account for genotype and sampling uncertainty. Such an approach facilitates obtaining probabilistic estimates of genetic variation that can be used to inform downstream parameters (Buerkle and Gompert 2013; Lou et al. 2021).

Another major concern with current genotype-environment association models is that they use phenomenological approaches, such as generalized linear regression models, to describe how evolutionary forces (e.g., selection, mutation, migration, and drift) interact to shape spatial genetic patterns (Goel et al. 2025). Although these phenomenological models are qualitatively consistent with evolutionary principles, they model genetic variation using statistical relationships that are often weakly motivated by evolutionary theory. Phenomenological statistical models may not account for important evolutionary processes, making it challenging to critique and improve them. For example, current genotype-environment association models account for migration mainly in the context of how it interacts with drift (Coop et al. 2010; Frichot et al. 2013): migration between demes produces spatial correlation in neutral allele frequencies that decays with distance, a pattern commonly referred to as isolation-by-distance (McRae 2006; Slatkin 1993). However, interactions between migration and selection at non-neutral loci are seldom accounted for in current methods, even though these interactions can be substantial when migration is structured and a species experiences steep environmental gradients (Hoekstra et al. 2004; Kirkpatrick and Barton 1997). One important consequence of the migration-selection balance is that a species may not exhibit a typical S-shaped response curve to environmental variation. Two locations experiencing high gene flow and divergent selection in response to the local environment can result in deviations from the S-shaped response curve. These deviations or irregularities stem from maladaptive gene flow, which cannot be easily accommodated by linear models.

A more effective approach is to use evolutionary theory to mechanistically model patterns of genetic variation directly (e.g., Goel et al. 2025). Because mechanistic models of evolution are constructed from first principles, we can evaluate statistical models based on their underlying biological assumptions (Rice 2004). More importantly, if necessary, we can refine the statistical model based on the species’ demographic history. This mechanistic approach to identifying adaptive loci has several advantages. The precise mathematical relationship between evolutionary parameters constrains the flow of information using scientific knowledge, rather than relying on heuristic functional forms (Hilborn and Mangel 2013). And because the parameters in an evolutionary model have biological meaning, the estimated parameters may provide valuable biological insights that are otherwise challenging to obtain using phenomenological models (Hooten and Hefley 2019; Wikle 2003).

In this paper, we developed a hierarchical Bayesian model (henceforth, referred to as the structured genotype-likelihood method) to address the aforementioned limitations. Our structured genotype-likelihood method consists of three levels: the data, process, and parameter models (Berliner 1996). The data model describes a probabilistic link between genetic data and genetic variation. The process model partitions this genetic variation into adaptive and non-adaptive components using a mechanistic evolutionary model. The parameter models represent our prior knowledge about the parameters. For example, neutral evolutionary theory suggests that most loci in the genome do not contribute to local adaptation (Kimura 1983) and, as such, should have a flat response curve.

The structured genotype-likelihood method has two key features. First, the method enables us to use genotype-likelihoods obtained from low-coverage whole-genome sequencing data to improve our statistical learning of genetic variation, which is then used to inform the parameters of a mechanistic evolutionary model in the downstream analysis. Second, our statistical approach relies on a new probabilistic metapopulation model of evolution to account for how structured migration (described using coalescent theory) interacts with local adaptation to modify the S-shaped response curve. To evaluate the robustness of our statistical method, we test and compare our statistical model against redundancy analysis, RDA (Capblancq and Forester 2021), and latent factor mixed models, LFMM (Caye et al. 2019), two commonly used genotype-environment association methods, across multiple synthetic datasets with known parameter values. We then analyzed genotype-likelihood data corresponding to 3.7 million SNPs to identify adaptive variation in rosy-finches (genus *Leucosticte*), a group of songbirds that predominantly breed in alpine environments.

### Rosy-finches

The radiation of North American rosy-finches is comprised of three main species: the Black Rosy-Finch, the Brown-capped Rosy-Finch, and the Gray-crowned Rosy-Finch. Together, the species complex spans western North America, ranging from Alaska, where they are found at sea level, to northern New Mexico, where they breed at elevations up to 4000 m (Funk et al. 2023). The majority of rosy-finch populations breed in tundra-like vegetation near and above the timberline (Fig. 1).

**Figure 1.**
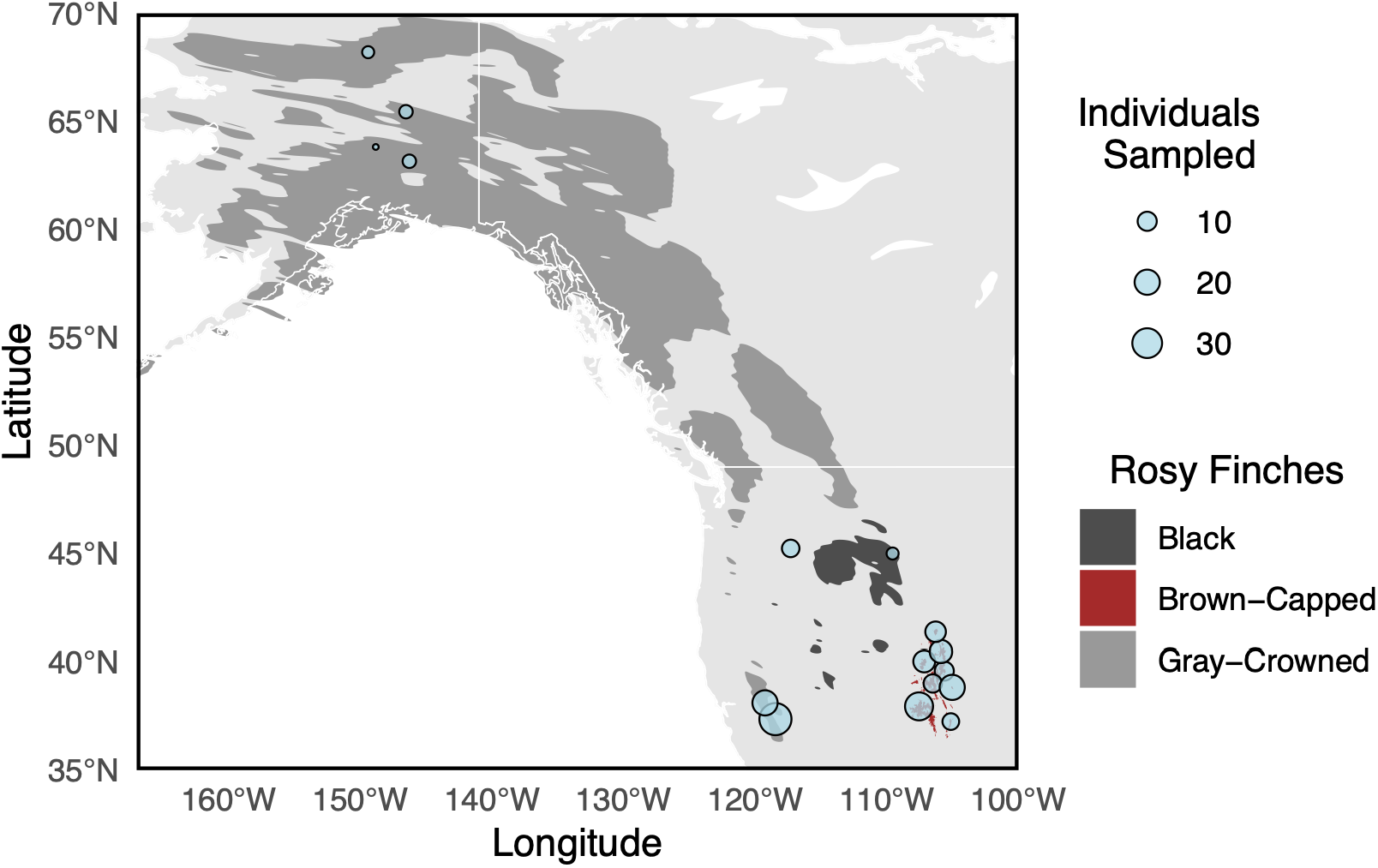
The breeding range of the North American rosy-finches spans the Western North America. The blue points denote the sampling sites, and the size of the points represents the number of birds sampled at the site. A total of 192 individuals were sampled. The sample sizes were highly heterogeneous, ranging from 1 to 36 individuals per sampling site.

## Data

A total of 192 individuals of rosy-finches were sampled at 24 sites across the United States. Because of the proximity of some sites, we merged those within 80 km of each other, yielding a total of 16 sites (Fig. 1, Table S1). Sample sizes were highly heterogeneous across sites, ranging from 1 to 36 individuals. Using a gradient forest method (Ellis et al. 2012), we identified the five least correlated environmental variables for our genotype-environment association analysis (see Table S2 for correlations between environmental variables): (a) mean July snowpack, (b) soil water content, (c) mean summer precipitation, (d) relative humidity, and (e) continentality. These environmental variables represent the three primary axes—snow persistence, seasonal moisture availability, and broad macroclimatic gradients—that structure the rosy-finch breeding habitat. July snow patches are key foraging sites during the breeding season when chick-provisioning is at its peak (Antor 1995; Brown et al. 2026). Soil water content is an integrative proxy for habitat quality in snowmelt microhabitat zones that support invertebrate prey base and seed-producing vegetation (Epanchin et al. 2010). Summer precipitation and relative humidity, together, capture seasonal moisture availability during the breeding period, which governs the reliability of snow-adjacent and wet-meadow foraging resources (Amirkhiz et al. 2025). Continentality captures the relative influence of oceanic and continental climate on temperature seasonality and precipitation regime. This climatic variable represents a broad-scale axis governing selective environments across the mainland range of North American rosy-finches (Funk et al. 2023). Please refer to the supplementary information for further details on the sampling and sequencing protocols.

## Model Formulation

In what follows, we describe our hierarchical Bayesian model for identifying loci that may contribute to local adaptation. Henceforth, we refer to this model as the structured genotype-likelihood method. To facilitate the interpretation of the statistical model, we provide a directed acyclic graph (Fig. 2) that describes how the model parameters are linked to each other and the genetic and environmental datasets.

**Figure 2.**
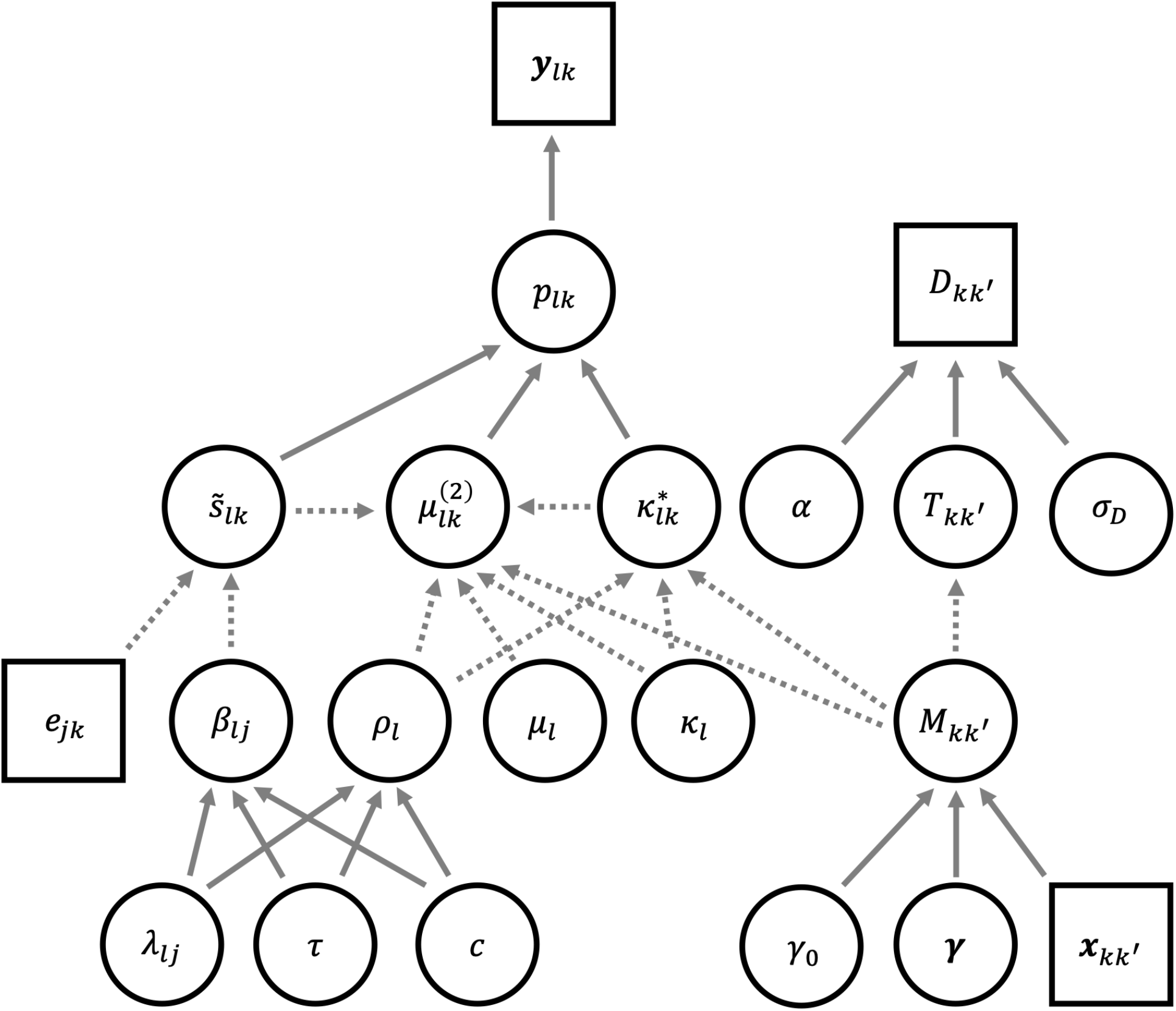
Directed acyclic graph (DAG) of the structured genotype-likelihood model. The squares represent data (reads and environmental variables), the circles represent parameters, the dashed arrows represent deterministic relationships, and the solid arrows represent stochastic relationships (see Eqs. 2, 9, 12, 14, and 17-26).

### Data model

We assume that individuals are sampled at *D* sites across a geographical region representative of the variation in the environment experienced by the species. A total of *N*_*k*_ individuals are sampled at site *k*. Each diploid individual is barcoded and sequenced at *L* polymorphic loci. However, due to finite sequencing depth, only *N*_*lk*_ individuals have read data corresponding to locus *l* (*N*_*k*_ ≥ *N*_*lk*_). We assume that the loci are biallelic and the genotype of an individual is an unknown parameter. To characterize the uncertainty in the genotypes of *N*_*lk*_ individuals, the read data (generically referred to as *y*_*ilk*_) for the *i*th individual are aligned to a reference genome. The alignment process yields genotype-likelihoods, 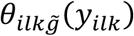, which corresponds to the probability that the genotype of the *i*th individual (*g*_*ilk*_) is equal to 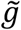 (*i*.*e*., 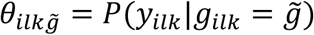). In this case, 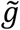 takes discrete values—zero, one, and two—and is equal to the number of copies of the reference allele. Intuitively, genotype-likelihoods, ***θ***_*ilk*_ = [*θ*_*ilk*0_, *θ*_*ilk*1_, *θ*_*ilk*2_]^*T*^, form a three-dimensional simplex vector, such that 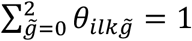. These genotype-likelihoods reflect uncertainty in the individual’s genotype arising from low-coverage, misalignment, and sequencing errors (Kim et al. 2011). To model genetic variation and coalescent time, we split the genetic dataset into two parts—*L*_1_ loci to model genetic variation and *L*_2_ loci to model coalescent time (*L*_1_ ≫ *L*_2_ and *L* = *L*_1_ + *L*_2_).

Assuming individuals are randomly sampled, we model the genetic variation at *L*_1_ loci as

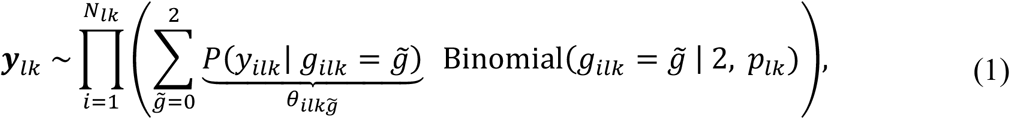

where 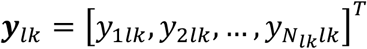 and *p*_*lk*_ are the read data and reference allele frequency in site *k* at locus *l*, respectively (Buerkle and Gompert 2013). Implicitly, the data model assumes conditional independence. We can re-express equation (1) as

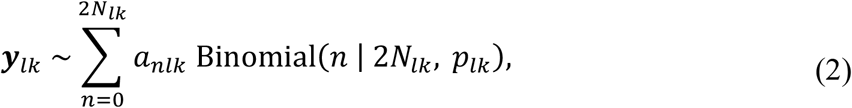

where the discrete parameter *n* denotes the total number of reference alleles in *N*_*lk*_ individuals. The Binomial distribution in equation (2) accounts for sampling uncertainty, and the scalar coefficients, *a*_*nlk*_, account for genotype uncertainty. These scalar coefficients are a function of 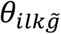 and *N*_*lk*_, and can be pre-computed using discrete convolution, such that

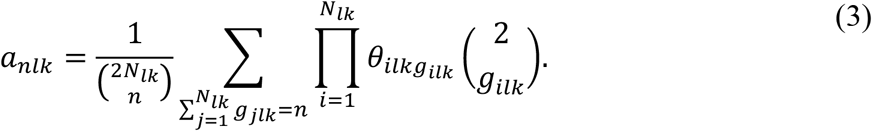

Note that equation (2) allows us to simultaneously account for uncertainty in the estimates of genetic variation arising from finite sampling and finite sequencing depth.

To quantify information content in the whole-genome sequencing data, we calculate the effective sample size (DeSaix et al. 2024)—the number of individuals with known genotypes that contain the same information about allele frequency as the genotype-likelihood data. More specifically, we calculate the number of sampled individuals with known genotypes and corresponding total reference allele counts that would yield the same posterior distribution of allele frequency as if one were to fit a standalone data model in equation (2) to genotype-likelihoods with *p*_*lk*_~Beta(1,1) prior. Using Bayes’ theorem, we can show that the posterior distribution of the allele frequency at locus *l* and site *k* is given by

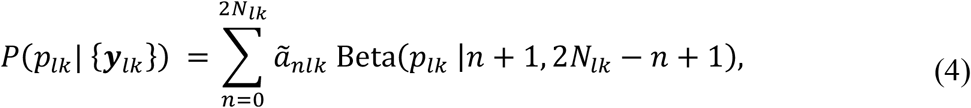

where 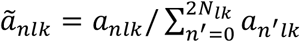. By moment matching, we can show that the effective sample size is

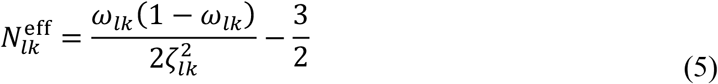

and the effective total counts of reference alleles is

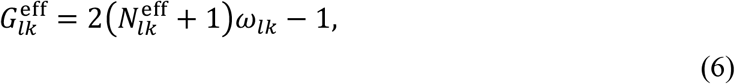

where

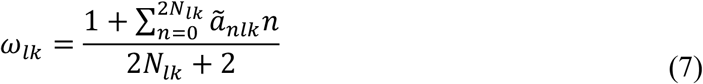

and

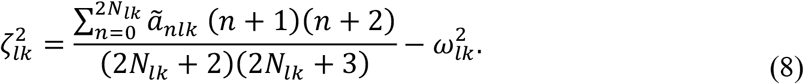

By defining effective sample size, 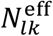, and the effective total counts of reference alleles, 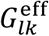, this way (Eqs. 5-8), we can faithfully capture the uncertainty in genetic variation that is embedded in the raw read data. For reference, Figure 3 shows how effective sample size varies with sequencing depth and the number of sequenced individuals.

**Figure 3.**
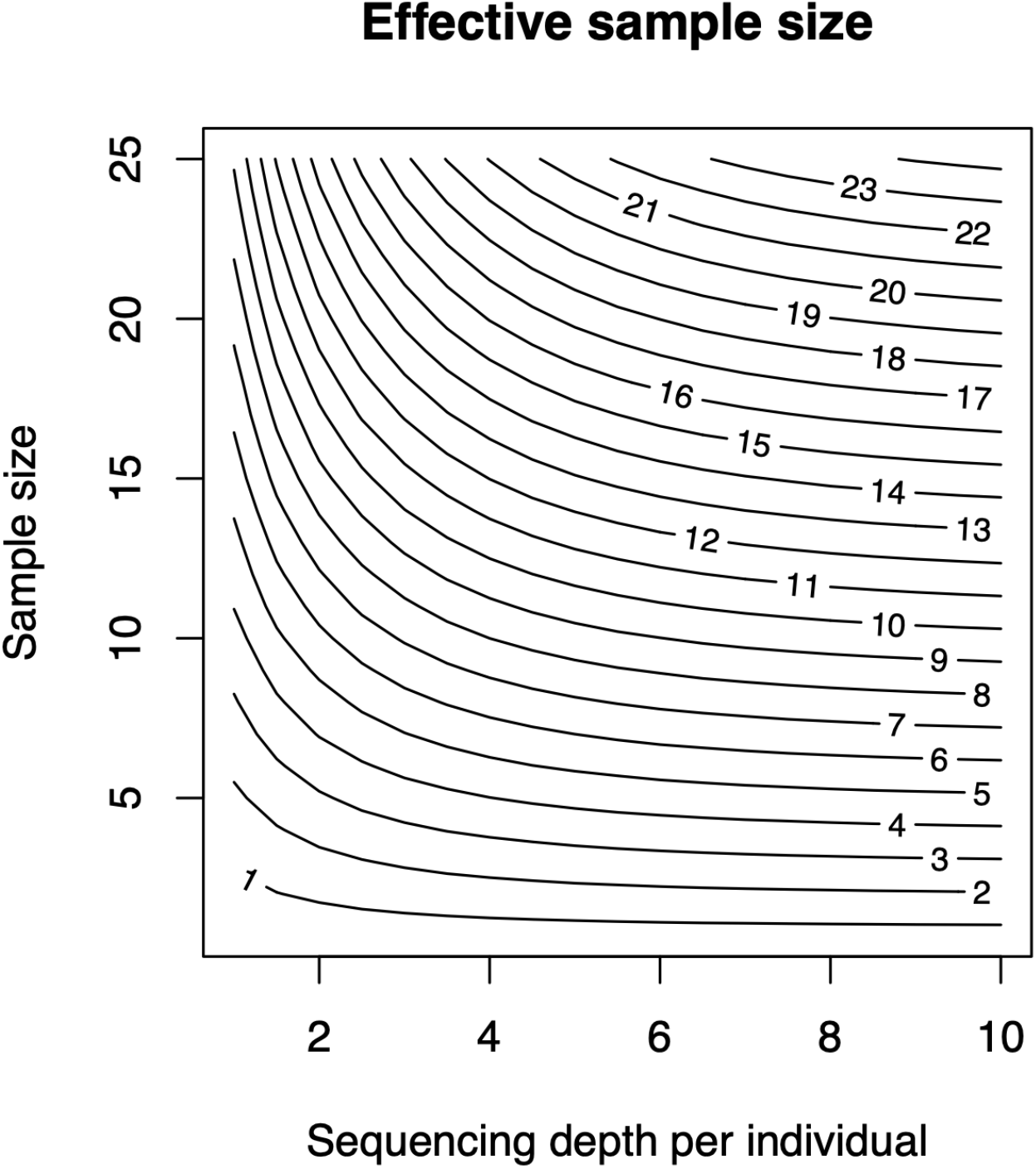
Effective sample size as a function of sequencing depth per individual and number of individuals sampled at a location. The contours represent combinations of sequencing depth and sample size for which the effective sample size is constant. Based on the desired level of precision in estimates of allele frequency (determined by the effective sample size), a biologist can make an informed decision about the relative allocation of resources between collecting individuals and sequencing effort.

To model coalescent times, we used *L*_2_ putatively neutral loci. Because the ‘true’ genotypes are unknown, we simulated *N*_sim_ = 50 genotype datasets (*g*_*ilk*_) using a categorical distribution with parameter ***θ***_*ilk*_ and categories zero, one, and two. For each dataset, we calculated per-deme reference allele frequencies 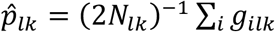. We then calculated within- and between-deme pairwise diversities 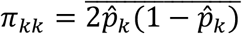 and 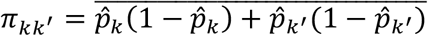 (averaged over *L*_2_ loci and *N*_sim_ samples). Following Petkova et al. (2016), the genetic dissimilarity between two diploid individuals is *D*_*kk*_′ = 4*π*_*kk*_′ − *π*_*kk*_′− *π*_*k*_′_*k*_′ and its expectation under neutral coalescent is proportional to 4*T*_*kk*_′ − *T*_*k*_′_*k*_′ − *T*_*kk*_′ (McVean 2009), where *T*_*kk*_′ is the expected coalescent time of two randomly chosen sequences from sampling site *k* and *k*^′^. We statistically model the observed genetic dissimilarity as

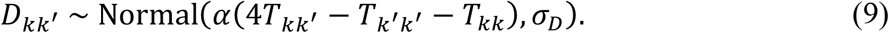

Although we formulate the data model for low-coverage whole-genome sequencing data, we can also specify data models for other types of commonly used genetic datasets, such as read data from sequencing whole genomes of pools of individuals (pool-seq for short, Gautier et al. 2013) and genotype data obtained from sequencing restriction site-associated DNA (RAD-seq for short, Baird et al. 2008) (see supplementary information for corresponding data models; Figs. S1-S3).

### Process model

We developed a structured metapopulation model to describe evolutionary dynamics, in which demes correspond to the sites where individuals were sampled. In each deme, the genetic variation is regulated by directional and non-directional evolutionary forces (Rice 2004). Directional forces, such as mutation, migration, and adaptation to local climate, yield a non-zero expected evolutionary change. In contrast, non-directional forces, such as genetic drift, yield stochastic evolutionary changes stemming from finite population size. We model these directional and non-directional evolutionary forces jointly to describe genetic variation at locus *l* using the following coupled stochastic differential equations (Korolev et al. 2010):

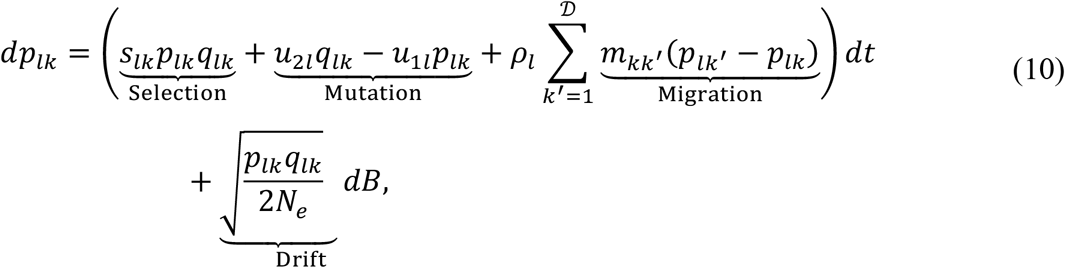

where *q*_*lk*_ = 1 − *p*_*lk*_ is the alternate allele frequency, *u*_2*l*_ and *u*_1*l*_ are the forward and backward mutation rates, *ρ*_*l*_*m*_*kk*_′ is the locus-specific symmetric migration rate between demes *k* and *k*^′^, *N*_*e*_ is the number of diploid individuals in each deme, and *s*_*lk*_ is the environmentally determined selection coefficient. To simplify our notation, we rescale time as *t* = 4*N*_*e*_*t*^′^, which yields the following equations of evolution:

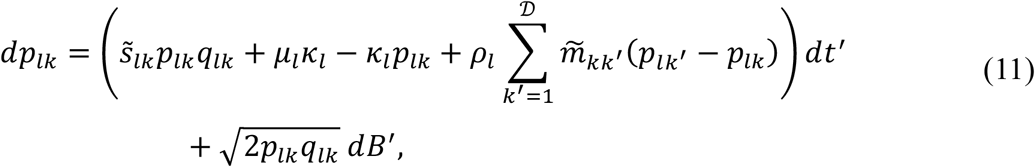

where 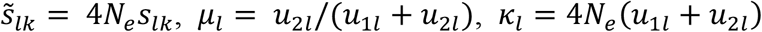, and 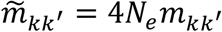. For simplicity, we use a linear relationship between (rescaled) selection coefficients and *E* environmental variables:

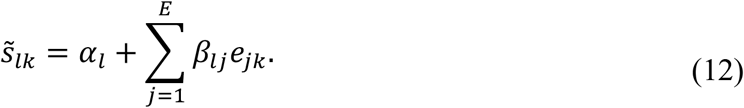

where *e*_*jk*_ is the *j*th environmental variable (standardized) in deme *k, β*_*lj*_ is the selection coefficient sensitivity to variation in the *j*th environmental variable, and *α*_*l*_ can be interpreted as the selection coefficient at mean environmental conditions (Goel et al. 2025).

The long-term evolutionary dynamics in equation (11) can be thought of as a balance between directional and non-directional evolutionary forces. The directional evolutionary forces drive the mean allele frequencies in the metapopulation to an equilibrium determined by selection, mutation, and migration. However, stochastic fluctuations in allele frequencies due to genetic drift result in deviations from the equilibrium allele frequencies. As the magnitude of the deviation increases, the directional evolutionary forces become stronger, pushing allele frequencies in the opposite direction and creating a range of probable values around the equilibrium. This range can be characterized by the stationary distribution, which is the solution of the multivariate Fokker-Planck equation corresponding to coupled stochastic differential equations that describe the evolutionary dynamics in equation (11) (Wright 1931).

We use this stationary distribution—the probability distribution of allele frequencies after a long time—as the process model to partition genetic variation into adaptive and neutral components (Goel et al. 2025). Although the exact solution of the multivariate Fokker-Planck equation is analytically intractable, we derive an approximation of the stationary distribution using an iterative substitution approach (Ortega and Rheinboldt 2000). First, we consider the baseline case in which migration is absent. In this case, the stationary distribution of the reference allele frequency is given by

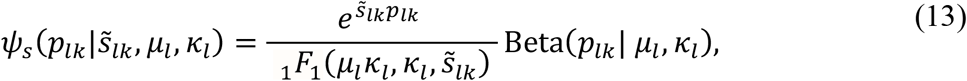

where _1_*F*_1_ is the confluent hypergeometric function (Wright 1931), and *μ*_*l*_ and *κ*_*l*_ are mean and precision parameters of the Beta distribution. The mean of this stationary distribution is given by

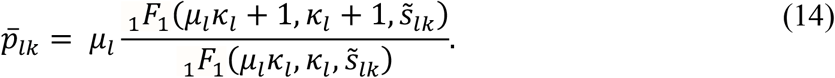

To include migration, we back-substitute 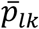 in equation (11), to get

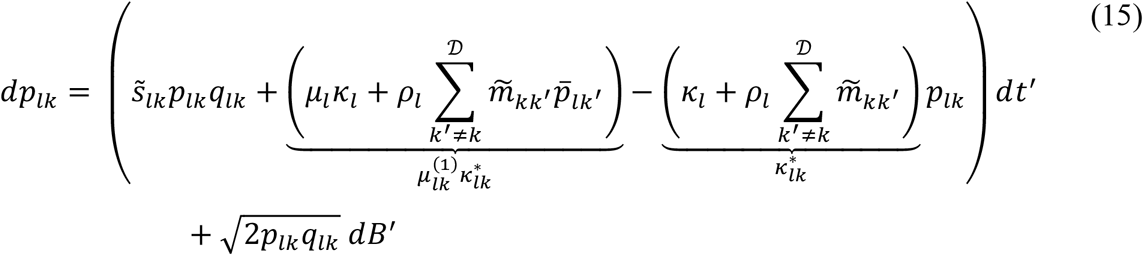

The substitution decouples the stochastic evolutionary dynamics of allele frequencies in demes, which allows us to analytically approximate the stationary distribution of allele frequencies under the joint influence of structured migration, local adaptation, mutation, and drift as

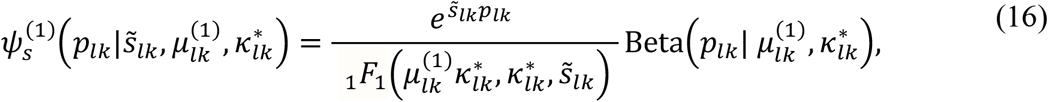

where

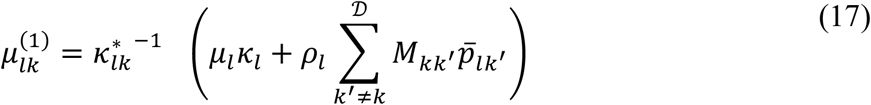

and

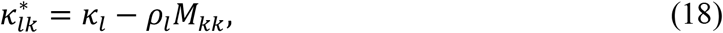

are the migration-corrected mean and precision parameters of the Beta distribution, respectively. We let ***M*** denote the symmetric migration-rate (generator) matrix, with 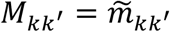 and 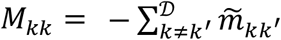. The migration-corrected mean of the stationary distribution, or response curve, is given by

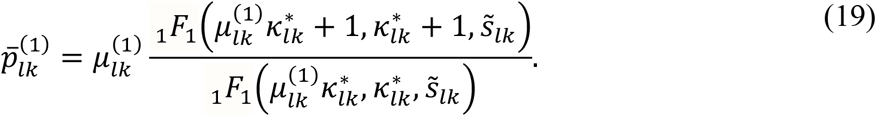

To further improve the approximation, we again substitute 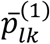 in equation (11) and re-calculate second-order migration-corrected 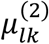, the stationary distribution 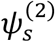, and the response curve 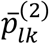. Our numerical simulations of the coupled stochastic differential equations (Eq. 11) confirm that 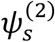 reasonably approximates the stationary distribution (see Fig. S4). More importantly, the migration-corrected response curve, 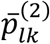, faithfully accounts for irregularities in the response curve resulting from maladaptive gene flow. Therefore, we use

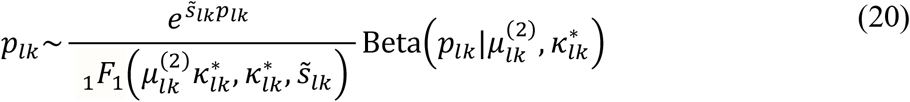

as our process model. Although higher-order iterative corrections to the stationary distribution (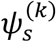 with *k* ≥ 3) may further improve the process model, our simulations show that these corrections offer only marginal improvements relative to the additional computational complexity.

Finally, we link the rescaled migration rates to coalescent times. Using the first-step analysis (Wakeley 2009), we can show that the coalescent time matrix (***T***) and migration matrix (***M***) are related as follows:

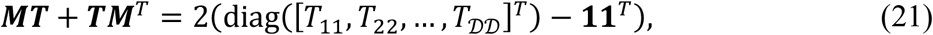

To constrain migration rates, we specify a dyadic regression to link migration rates between demes to the intervening features of the landscape (Schwob et al. 2024):

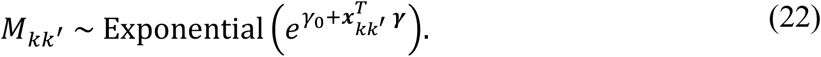

We let ***γ***, *γ*_0_, and ***x***_*kk*_′ denote the regression coefficients, intercept, and dyadic covariates integrated over the least-cost path from deme *k* to deme *k*^′^, respectively (see supplementary information for details on the covariates).

### Parameter model

We specify relatively diffuse priors for parameters that respect their natural bounds: *μ*_*l*_~Uniform(0,1), *κ*_*l*_~Normal^+^(0,4), ***γ***~Normal(**0, *I***), *γ*_0_~ Normal(0,1), *α*~Exponential(1), and *σ*_0_~Exponential(1). Due to the degeneracy of the likelihood (Eq. 20), the mean of the Beta distribution in equation (20) 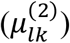 and selection coefficient 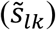 are non-identifiable parameters (Goel et al. 2025). To address this issue, we first fix *α*_*l*_ = 0. Next, based on our prior knowledge that only a small fraction of loci likely confer an adaptive advantage to the species, we use regularized horseshoe priors for sensitivity coefficients (*β*_*lj*_) (Goel et al. 2025; Piironen and Vehtari 2017). The regularization is controlled by a global shrinkage parameter, *τ*, which shrinks all sensitivity coefficients to zero, and a heavy-tailed local shrinkage parameter, 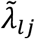, that allows a small fraction of sensitivity coefficients to take large values. The cumulative effect of these shrinkage parameters can be expressed as

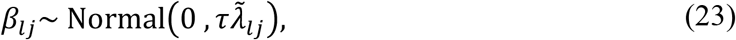

where

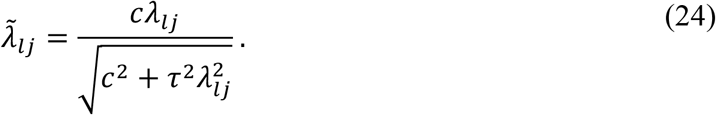

To control global shrinkage, we specify a half-Cauchy prior,

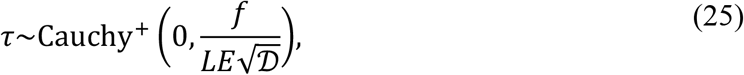

where *f* is a free variable. Based on the numerical experiments by Goel et al. (2025), we set *f* to 20. For the local shrinkage parameter, we use a heavy-tailed prior *λ*_*lj*_~Cauchy^+^(0,1) to allow some sensitivity coefficients to escape shrinkage. However, these large-valued coefficients are also weakly shrunk with an implied regularization of Normal(0, *c*) with *c*^2^ ~ InvGamma(2,4). We adopt an inverse-gamma prior because the prior mean for *c*^2^ (which controls adaptive dynamics via sensitivity coefficients) is similar in magnitude to prior means of *κ*_*l*_ (which govern neutral dynamics). However, the heavy right tail of the inverse-gamma distribution allows adaptive loci to attain large sensitivity coefficients. These features of the local and global shrinkage parameters effectively induce a spike-and-slab prior (Mitchell and Beauchamp 1988), which prevents numerical pathologies when large-effect loci are weakly identifiable. Finally, we model *ρ*_*l*_ as

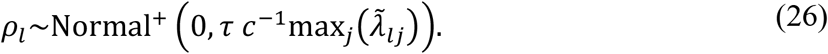

Intuitively, equation (26) creates a switch mechanism: for neutral loci *ρ*_*l*_ → 0 and for adaptive loci *ρ*_*l*_ has an implied distribution of Normal^+^(0,1). Our numerical experiments show that this switch mechanism considerably reduces overfitting and false positives.

### Improving sampling efficiency

Our initial model-fitting experiments revealed that the above specification of the structured genotype-likelihood model requires a considerable amount of memory and time to fit, even for reasonably sized datasets. To address these challenges, we first compute relative migration rates (***M***) using the model of genetic dissimilarity (Eq. 9), coalescent equations (Eq. 21), and dyadic regression (Eq. 22):

*P*(*α, σ*_*D*_, ***T, γ***, *γ*_0_|{*D*_*kk*_′})

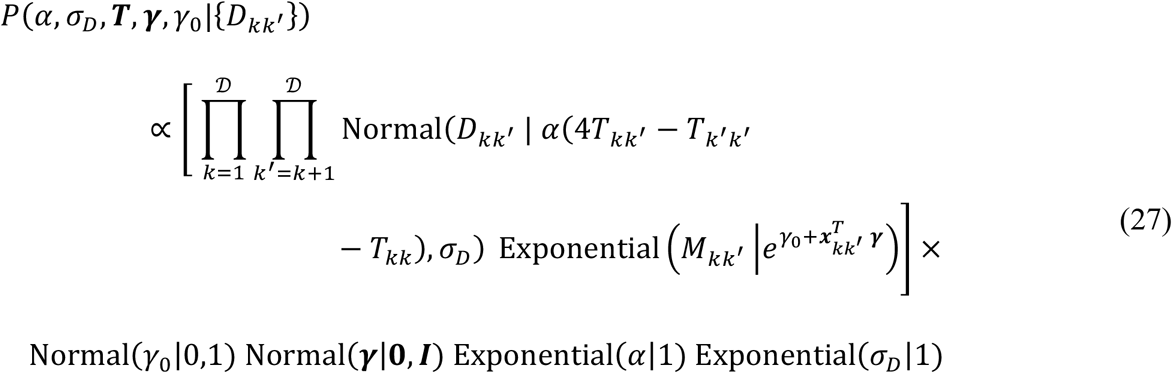

Normal(*γ*_0_|0,1) Normal(***γ***|**0, *I***) Exponential(*α*|1) Exponential(*σ*_*D*_|1)

We then treat mean estimates of migration rates as known inputs in the Bayesian model, while acknowledging that conditioning on them may lead our statistical model to underestimate parameter uncertainty.

Next, we consider the implied data model for the effective total counts of reference alleles

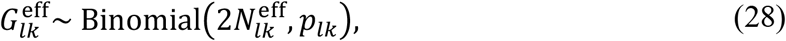

which approximately contains the same information about the allele frequencies as the original data model in equation (2). However, the new data model allows us to marginalize over *p*_*lk*_ and the total number of reference alleles (*n*), which substantially reduces the computational complexity. The marginalization yields the following joint moment-matched likelihood:

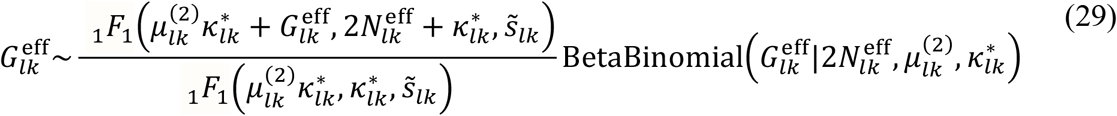

where the 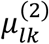 and 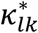 are the migration-corrected mean and precision parameters of the Beta-binomial distribution. This simplification reduces memory usage, improves numerical stability, and increases implementation speed, while accommodating the uncertainty inherent in the stochastic nature of sampling, sequencing, and evolutionary dynamics. Cumulatively, our streamlined data, process, and parameter models yield the following posterior distribution:

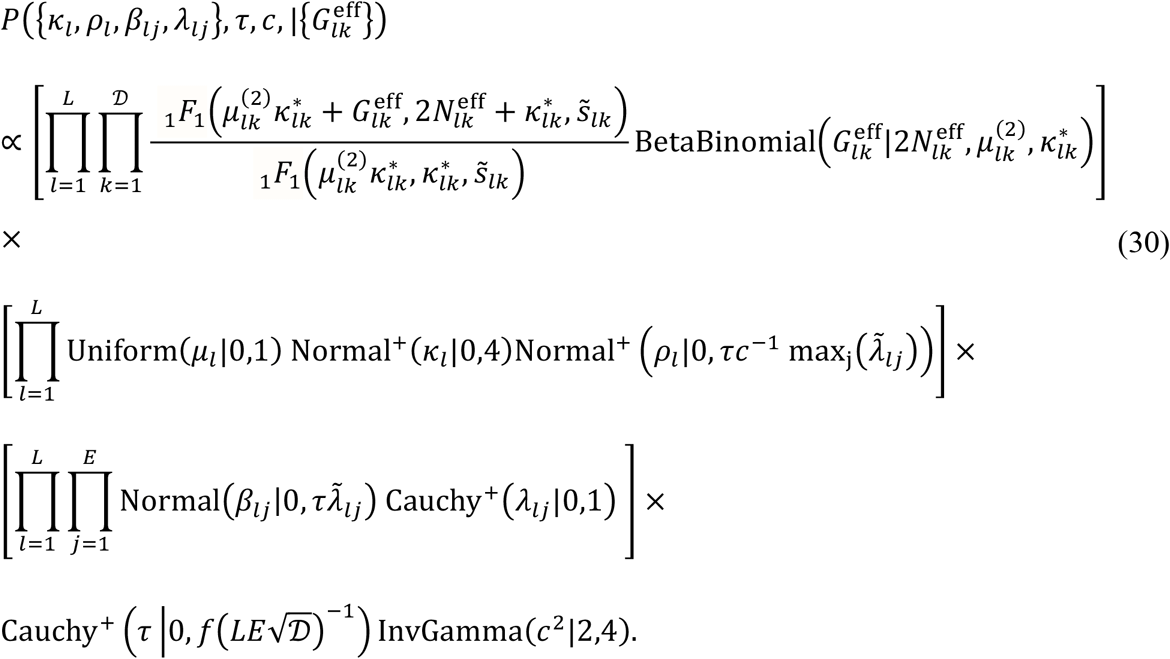

Our numerical experiments showed that pre-computing migration rates and marginalization reduce the fitting time by an order of magnitude, without qualitatively changing the inferences (see Model Testing section below).

## Model Testing

To test the robustness of our statistical model, we analyzed multiple synthetic datasets. We considered 5 migration regimes (low to high), 5 sequencing depths (low to high), and 5 iterations, yielding a total of 125 synthetic datasets. For each combination of iteration, sequencing depth, and migration regime, we generated the synthetic dataset in three stages. First, we randomly chose 20 coordinates in a 2D landscape, which we treat as demes. Migration between demes decreases exponentially with distance. A deme is characterized by 5 environmental covariates, each drawn from a standard normal distribution. At each deme, we sampled 10 diploid individuals, each with a genome of 2000 independent loci. We used 500 loci (*L*_1_) to fit the 6 GEA methods and 1500 loci (*L*_2_) to estimate migration rates for the structured genotype-likelihood and structured called-genotype methods. We assigned adaptive loci a small value of *α*_*l*_ drawn from Normal(0,0.5). We chose three loci per environmental variable and assigned them large sensitivity coefficients (|*β*_*lj*_| = 15), with their signs determined randomly. For all *L*_2_ loci, we set the selection coefficient to zero.

In the second stage, we simulated allele frequencies, *p*_*lk*_, using the stochastic equations of evolution (Eq. 10) with selection coefficients from the first stage as input. In the final stage, we simulated genotype-likelihoods (***θ***_*ilk*_) for individuals at a given sequencing depth. For the *i*th individual at locus *l* in deme *k*, we sampled its genotype (*g*_*ilk*_) from a Binomial(2, *p*_*lk*_) distribution, where *p*_*lk*_ is obtained from stage two of the simulations. Then, we drew a random variable from the Poisson distribution with mean equal to sequencing depth to represent the total number of sequencing reads at locus *l* (*n*_*ilk*_). To generate read counts corresponding to the reference allele 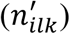, we sampled from a Binomial(*n*_*ilk*_, 0.5*g*_*ilk*_(1 − *ϵ*) + (1 − 0.5*g*_*ilk*_)*ϵ*) distribution, where *ϵ* (= 0.005) is the probability of sequencing error. Finally, we generated genotype-likelihoods using *ϵ*, 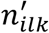, and *n*_*ilk*_ as inputs in equation (3) in Buerkle and Gompert (2013). To generate called genotypes, we assigned each individual the genotype corresponding to the highest genotype-likelihood.

For each dataset, we tested 6 genotype-environment association methods: (*i*) structured model presented in this paper with genotype-likelihoods as input data, (*ii*) unstructured model with genotype-likelihoods as input data (adapted from Goel et al. 2025), (*iii*) unstructured model with called genotypes as input data (adapted from Goel et al. 2025), (*iv*) structured model presented in this paper with called genotypes as input data, (*v*) RDA with called genotypes as input data (Capblancq and Forester 2021), and (*vi*) LFMM with called genotypes as input data (Caye et al. 2019). The comparison between methods (*i*), (*ii*), (*iii*) and (*iv*) allowed us to evaluate whether including migration and genotype uncertainty improves inferences. The comparison between methods (*iv*), (*v*), and (*vi*) helped us evaluate the performance of our model against commonly used GEA methods. Note that for the second comparison, we did not use the structured genotype-likelihood method. Instead, we used the structured called-genotype method because LFMM and RDA do not accept genotype-likelihoods as inputs.

For the structured called-genotype method, we used the RAD-seq data model presented in the supplementary information. For the unstructured genotype-likelihood method, we used effective sample sizes and effective total counts of reference alleles as data in equation (13) in Goel et al. (2025). For the unstructured called-genotype method, we used called genotypes as data in equation (13) in Goel et al. (2025). The performance of each method was evaluated by counting false positives and false negatives. The first four models were implemented in Stan (Carpenter et al. 2017), and LFMM and RDA were implemented using the *lfmm* and *vegan* packages in the R programming language, respectively (Capblancq et al. 2018; Caye et al. 2019).

Across all synthetic data sets, the structured genotype-likelihood method presented in this paper yielded the fewest false negatives compared to the unstructured genotype-likelihood and structured called-genotype methods, albeit with higher false positives (see Table 1). As expected, the structured genotype-likelihood method performed best in high migration and low sequencing-depth regimes, underscoring the importance of accounting for genotype uncertainty and structured migration for identifying adaptive variation (Fig. S5). Furthermore, the structured called-genotype method also yielded lower false negatives (but slightly higher false positives) when compared with LFMM and RDA with called genotypes as inputs (Table 1 and Fig. S6).

**Table 1.** False negative (column 4) and false positive (column 5) rates for 6 genotype-environment association methods (column 1) across all synthetic datasets. These methods include (*i*) structured model presented in this paper with genotype-likelihoods as input data, (*ii*) unstructured model with genotype-likelihoods as input data (adapted from Goel et al. 2025), (*iii*) unstructured model with called genotypes as input data (adapted from Goel et al. 2025), (*iv*) structured model presented in this paper with called genotypes as input data, (*v*) RDA with called genotypes as input data (Capblancq and Forester 2021), and (*vi*) LFMM with called genotypes as input data (Caye et al. 2019). Also see Figures S5 and S6 for a comparison of synthetic datasets across different combinations of migration rates and sequencing depth.

| Method | Input data | Citation | False negative % | False positive % |
| --- | --- | --- | --- | --- |
| Structured | Genotype-likelihoods | This paper | 6.99 | 0.475 |
| Unstructured | Genotype-likelihoods | Adapted from Goel et al. (2025) | 67.41 | 0.001 |
| Unstructured | Called genotypes | Adapted from Goel et al. (2025) | 67.09 | 0.002 |
| Structured | Called genotypes | This paper | 18.08 | 0.128 |
| RDA | Called genotypes | Capblancq and Forester (2021) | 33.12 | 0.034 |
| LFMM | Called genotypes | Caye et al. (2019) | 31.36 | 0.073 |

For illustration, the Manhattan plot in Fig. 4A shows the results from the structured genotype-likelihood method presented in this paper for medium migration and a sequencing depth of 4. To plot the results, we obtained the posterior samples corresponding to sensitivity coefficients (*β*_*lj*_) for *L*_1_ loci. For each pair of loci and environmental variables, we computed the probability that the posterior distribution of the sensitivity coefficient included zero. If this probability was less than the threshold value 0.05 (*p*_th_), we inferred that the locus could have contributed to local adaptation to the corresponding environmental variable. In Fig. 4A, the y-axis represents the negative log probability that the posterior distribution of sensitivity coefficients included zero (Wang et al. 2022b). Thus, the points above *y* = 2.99 (the black line) have *p*_th_ < 0.05. These points are characterized by a ring and a solid center. The color of the center represents the true environmental variable associated with the loci, and the color of the ring represents the statistically inferred adaptive environmental variable. We also re-analyzed the data in Fig. 4A by fitting the full Bayesian model with migration rates (***M***) and total number of reference alleles (*n*) as parameters (see supplementary information for the full Bayesian likelihood). Our results were similar to the results in Fig. 4A (see Fig. S7).

**Figure 4.**
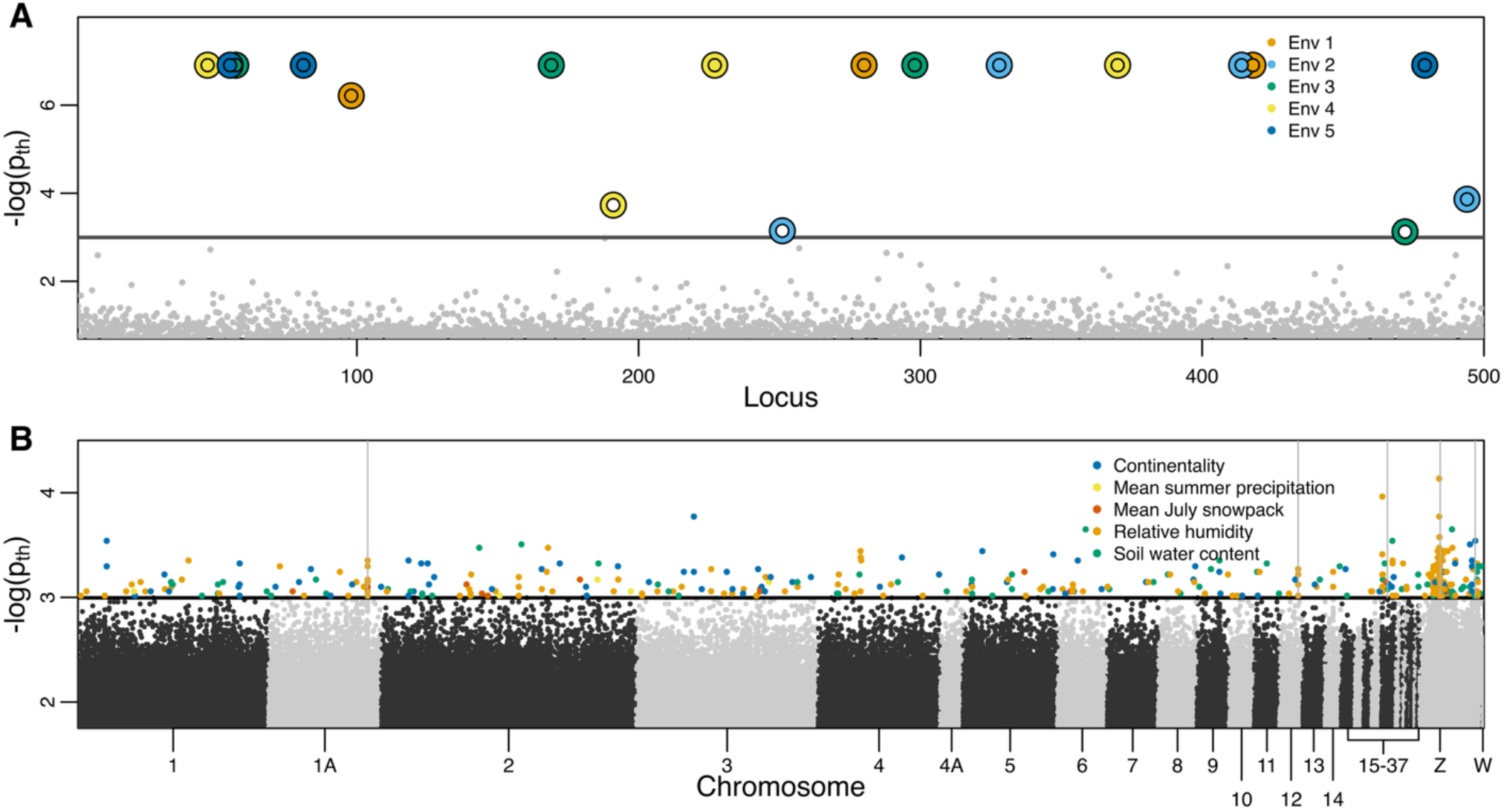
Manhattan plot showing negative log probability (− log(*p*_th_)) that the posterior distribution of sensitivity coefficients (*β*_*lj*_) includes zero for synthetic (A) and rosy-finch data (B). In both plots, points above the black horizontal line have *p*_th_ less than 0.05. (A) For synthetic data, we denote points above the black line using an outer ring and a center. The color of the center (ring) corresponds to the true (statistically inferred) environmental variable responsible for local adaptation. (B) For real data, we denote points above the black line with a solid center, with its color corresponding to the statistically inferred environmental variable responsible for local adaptation. The vertical gray lines represent physically linked loci with statistically significant sensitivity coefficients.

## Data Analysis

We identified a total of 3.87 million high-quality variants after filtering for quality, missingness, and batch effects. After aligning the reads with the Zebra Finch reference genome, we recovered approximately 3.75 million SNPs. We used 50 thousand SNPs to estimate mean relative migration rates (Eq. 27), which were subsequently used as known inputs in the Bayesian model (Eq. 30). Computing migration rates required 2 minutes. To analyze the remaining 3.7 million SNPs, we partitioned the genotype-likelihood data into 37 non-overlapping 100k SNP batches and analyzed each batch concurrently on ICX compute nodes on Stampede3, Texas Advanced Computing Center (Boerner et al. 2023). Each ICX node features 80 cores with a nominal clock rate of 2.3 GHz (max frequency: 3.4 GHz) and 256 GB of RAM. Our simulations took 16 hours (or cumulatively 25 days of compute time) to fit the Bayesian model (Eq. 30). We also reimplemented our model in JAX (Bradbury et al. 2018), with sampling in NumPyro’s NUTS (Phan et al. 2019), and it took ~20 minutes to fit 100k SNPs (or cumulatively ~12 hours of compute time) on an H100 compute node on Stampede3. Using one of the SNP batches, we also computed the effective sample size.

Our data simulations revealed that the genotype-likelihood data resulted in an effective sample size of 69% of the sampled birds. Our statistical method identified 434 candidate loci with statistically significant sensitivity coefficients. We found that 57% of these outlier loci were associated with relative humidity, 18% with soil water content, 21% with continentality, 3% with the average July snowpack, and 1% with mean summer precipitation (Fig. 4B). About 48% of the statistically significant loci were clustered together on the genome due to genetic hitchhiking (gray vertical lines in Fig. 4B).

Of the 434 loci, 347 were located within 25 kb upstream or downstream of 231 named genes, which is thought to be the distance within which linkage disequilibrium is expected to break down (Backström et al. 2006). In addition, we compiled a list of candidate genes from the literature with gene ontology (GO) terms that could be linked to an alpine specialist, for example, associated with aspects of thermal tolerance, as well as genes linked to avian thermal stress and/or bill morphology. We also included genes previously identified in the rosy-finch species complex (Funk et al. 2021; Funk et al. 2023), as well as migratory-linked candidate genes previously identified in other species. We then searched for genes within 25 kb of these candidate loci in the published candidate list and for genes with relevant GO terms to examine the potential functions of our identified candidate SNPs and further narrowed our focus to 28 candidate genes in close proximity to statistically significant loci (Table S3). These genes broadly fall into 5 functional categories: pigmentation (7 genes), migration (5 genes), physiology linked to high-altitude adaptation (8 genes), physiology linked to stress response (3 genes), and beak and feather development (5 genes) (see Table S3).

The relationship between genetic dissimilarity and distance between demes suggests that North American rosy-finches do not show an isolation-by-distance pattern (Fig. 5A). In particular, we found that the Gray-crowned Rosy-Finches in the Sierra Nevada, California, exhibit low connectivity with the rest of the species complex (Fig. 5B). Our results show that migration-selection balance can play an important role in shaping genetic variation (Fig. 6). When migration is absent, allele frequencies (represented by gray squares in Fig. 6) follow a typical S-shaped response curve. However, when migration is present, we observed that allele frequencies (represented by orange points in Fig. 6) regress to the mean, and the response curve exhibits irregularities.

**Figure 5.**
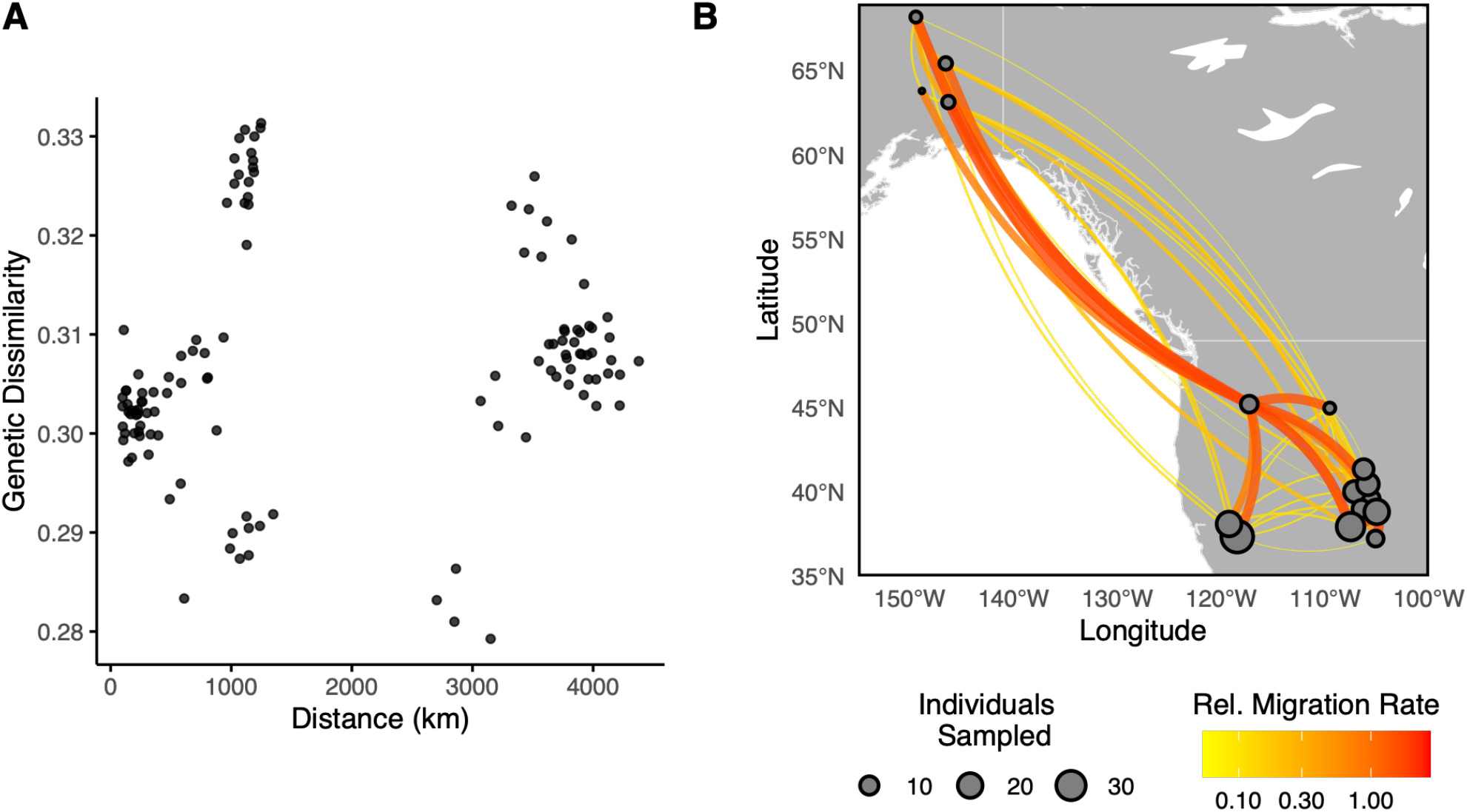
Genetic dissimilarity (A) and migration map (B) of North American rosy-finches. The genetic dissimilarity of rosy-finches does not increase with geographical distance between demes, suggesting heterogeneous landscape connectivity.

**Figure 6.**
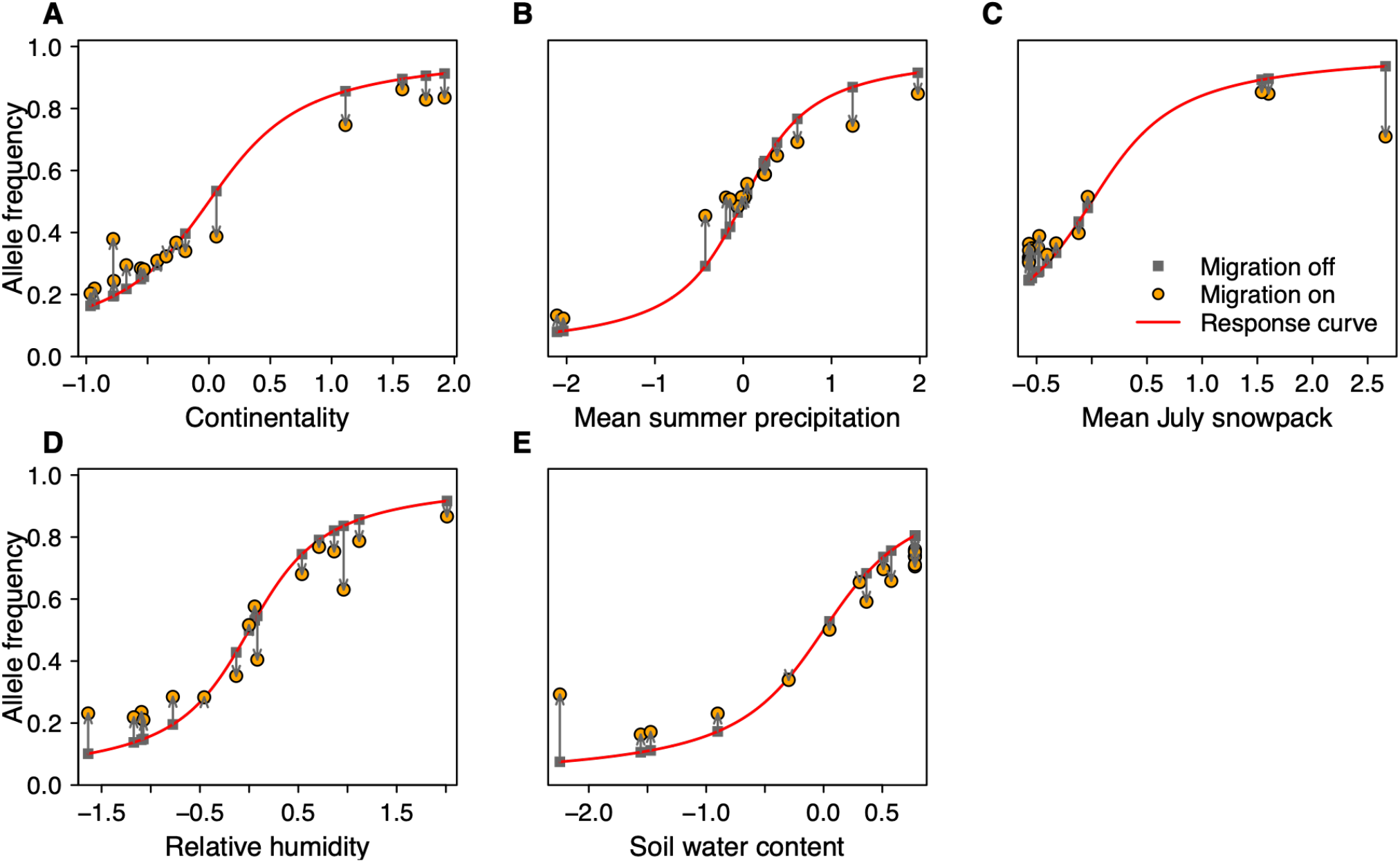
Relationship between allele frequency and environment when migration is present (orange points) and absent (gray squares). In the absence of migration, the response curve, 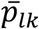, takes a typical S-shape curve (red line). However, when we account for migration, we observe irregularities in the response curve 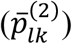, suggesting that migration-selection balance can lead to non-clinal patterns in allele frequencies. To generate these plots, we use migration rates inferred from rosy-finch genetic data (Fig. 5). For all plots, we use parameter values *μ*_*l*_ = 0.5, *κ*_*l*_ = 3.5, *ρ*_*l*_ = 1, and *β*_*lj*_ = 10.

## Discussion

We presented a highly parallelizable hierarchical Bayesian model to identify genomic adaptation in spatially structured populations (Fig. 2 and Eq. 30). Our statistical model extends the previous approach of Goel et al. (2025) and other currently available genotype-environment association methods (Rellstab et al. 2015) in two respects. We present (a) a probabilistic data model (Eqs. 1-8 and 28) to estimate genetic variation using low-coverage whole-genome sequencing data, and (b) a novel process model of evolution (Eq. 20) to account for migration-selection balance when fitting a clinal relationship between allele frequency and environmental covariates 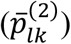. Our approach is one of the first methods for identifying adaptive loci under selection from the environment while accounting for migration.

### Data model for low-coverage whole-genome sequencing data

To model low-coverage whole-genome sequencing data, we propose a binomial data model (Eq. 28). This binomial model uses effective sample size and genotype counts (Eqs. 4-8) as inputs, allowing us to simultaneously capture sampling and genotype uncertainty arising from finite sampling and sequencing effort, respectively. This methodological approach has two advantages (Lou et al. 2021). First, accounting for sampling uncertainty enables us to include sites with extremely low sample sizes (1-3 individuals). Even though these sites individually may not contain much information about underlying genetic variation in the landscape, collectively they may contain non-trivial information to constrain the shape of the response curve, resulting in more robust statistical inferences. Second, by explicitly accounting for and propagating genotype uncertainty, our statistical model avoids systematic biases from false genotype calls and provides more precise estimates of genetic variation across a broader genomic region (see the comparison between structured genotype-likelihood and structured called-genotype methods in Table 1 and Fig. S5).

Accounting for genotype uncertainty also offers opportunities to design more affordable sampling and sequencing protocols by spreading the sequencing effort over a larger number of individuals (Buerkle and Gompert 2013; Lou et al. 2021). As an example, our simulations showed that sequencing 20 individuals at 2x (genotypes cannot be called reliably and the total sequencing effort is 40x) yields the same information about allele frequency as sequencing 12 individuals at 30x (genotypes can be called reliably and the total sequencing effort is 360x), even though the two strategies differ by an order of magnitude in total sequencing effort.

To aid practitioners in their study design, we show in Figure 3 how effective sample size— which determines the precision of estimates of allele frequency—varies with sequencing and sampling effort. The figure can be used to determine how to distribute resources between sequencing and sampling to achieve a desired effective sample size. When human resources are limited or the species occurs in remote areas, one can devote more resources toward sequencing. Alternatively, for species that are easy to collect, such as many plants, one can benefit by collecting a larger sample size. Surprisingly, using low-coverage whole-genome sequencing data does not incur additional computational cost compared with using traditional called-genotype data, because effective sample size and genotype counts can be evaluated cheaply. For the rosy-finches dataset, it took under 30 minutes to calculate effective sample sizes and effective total counts of reference alleles, which is substantially less than 25 days of compute time to fit the structured genotype-likelihood method.

### A mechanistic structured metapopulation model of evolution

Another main feature of our statistical method is a new mechanistic process model of evolution (Eqs. 16-20). This process model corresponds to the stationary distribution of allele frequencies shaped by the joint interactions between local adaptation, mutation, structured migration, and drift. We derive an approximate stationary distribution (Eq. 20) corresponding to the stochastic evolutionary dynamics in equation (11). A key feature of our process model is that it allows us to mechanistically account for the interactions between selection (measured by sensitivity coefficients; see Eq. 12) and migration by learning these parameters from genetic data using regularization (Hooten and Hobbs 2015; Eqs. 23-25) and structured coalescent theory (Eq. 21), respectively. These interactions can be substantive when species movement is structured, and the landscape has steep environmental gradients.

Our analysis of rosy-finch genomes and synthetic genetic datasets reveals that accounting for migration can provide new biological insights. Estimates of migration rates suggest that rosy-finches do not exhibit isolation by distance, a commonly observed pattern when migration is distance-dependent (Slatkin 1993) (Fig. 5A). Instead, we find strong connectivity between distant locations (Alaska and populations in the Western United States) and weak connectivity between nearby locations (Sierra Nevada and other populations in the Western United States), indicating heterogeneous migration rates (Fig. 5B). The observed patterns of population connectivity could be related to the diverse non-breeding season movement behavior of North American rosy-finches. Specifically, populations that breed in Alaska and Canada are known to make long-distance movements to the Western United States during the non-breeding season and form mixed-species flocks with other species of rosy-finches and other subspecies of Gray-crowned rosy-finches (Campbell et al. 2025; MacDougall-Shackleton et al. 2000). Because pair formation is thought to occur late in the non-breeding season (MacDougall-Shackleton et al. 2000), such population mixing during the non-breeding season could facilitate gene flow between spatially distant breeding populations. The Sierra Nevada population, on the other hand, is thought to exhibit short-distance facultative movements to lower elevation areas east of the Sierra Nevada (MacDougall-Shackleton et al. 2000), which likely provide fewer opportunities for mixing between distant breeding populations during the non-breeding season.

Accounting for migration can also increase the statistical power for identifying adaptive variation in structured populations (see the comparison between structured genotype-likelihood and unstructured genotype-likelihood methods in Table 1 and Fig. S5) because of the better approximate stationary distribution (see Fig. S4). In simple cases, such as when migration between patches is either absent or global, theory predicts a smooth S-shaped response curve (gray points in Fig. 6). However, the distributions of most species are spatially structured, with complex migration patterns (e.g., Fig. 5). In this realistic scenario, the migration-selection balance can introduce irregularities in the response curve (orange points in Fig. 6) due to maladaptive gene flow from environmentally distinct locations. Consequently, statistical methods that do not account for structured migration will lose substantial power when fitting a smooth S-shaped response curve to allele frequencies that do not vary smoothly with environmental covariates, potentially leading to higher false-negative rates. To address this challenge, our method allows us to separate the irregularities due to structured migration from the observed response curve before fitting a smooth S-shaped function 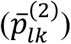.

### Analysis of synthetic and real data

Synthetic simulations show that the structured called-genotype method yields lower false negatives than LFMM and RDA in high-migration and low sequencing-depth regimes (Table 1 and Fig. S6). Furthermore, analysis of 3.7 million SNPs from rosy-finch genomes revealed 434 loci putatively under selection, of which 48% were clustered together due to genetic hitchhiking (gray vertical lines in Fig. 4B). Gene ontology revealed 28 candidate genes linked to pigmentation, migration, physiology linked to high-altitude adaptation, physiology linked to stress response, and beak and feather development (Table S3). Our method identified eight genes previously documented in rosy-finch genomic analyses, even after adding new differentiated populations and individuals (Funk et al. 2023; Funk and Taylor 2019). Four of the genes linked to high-altitude adaptation (AGGF1, ALDH1A1, *EGLN1*, and JMY) were previously found by Funk et al. (2023). Here, we identified two additional genes with known links to adaptability in hypoxic high-altitude environments, *EPAS1 and TNFSF11*, and two associated with thermal tolerance in extreme environments, *PPARGC1A* and *RTEL1*. Notably, *EPAS1* forms a part of the HIF (hypoxia-inducible factor) complex that enhances oxygen delivery in hypoxic environments (Beall et al. 2010; Graham and McCracken 2019; Simonson et al. 2012; Yi et al. 2010). *TNFSF11*, also known as RANKL, is enhanced by HIF1α in simulated hypoxic environments and promotes osteoclast differentiation and bone resorption (Wang et al. 2022a). The latter two genes are associated with extreme environments: *PPARGC1A* with cold tolerance and water scarcity in chickens (Gheyas et al. 2021) and *RTEL1* with body temperature changes when subjected to acute heat (Zhuang et al. 2019). Moreover, *RTEL1* variants were associated with the risk of high-altitude pulmonary edema in the Chinese Han population (Rong et al. 2017). We also identified two genes that may be important for adaptation to the alpine environment—one that regulates the stress response (CRHBP; Wan et al. 2022) and another that protects cells from oxidative stress (GPX8; Aryal et al. 2025; Pei et al. 2023). Five of seven pigmentation genes are associated with the melanin pathway, which may relate to known differences in the extent of dark coloration on the head and crown across the species complex. Further, melanin pigmentation differences may also shield against intense solar radiation at different elevations in line with Gloger’s rule (Mayr 1942; Rensch 1938).

### Future Improvements

The capabilities of our statistical method can be further expanded by integrating alternate data models to analyze called-genotype data obtained by RAD sequencing and read data from pool sequencing (see supplementary information for a brief description of how to integrate data model for RAD- and pool-seq data with the mechanistic process model in Eq. 20). To analyze RAD-seq data (Baird et al. 2008), one can calculate the total number of reference alleles in the sampled individuals and model the allele frequency as a binomial distribution to account for sampling uncertainty (Coop et al. 2010; De Villemereuil and Gaggiotti 2015; Goel et al. 2025) (Fig. S1).

To analyze pool-seq data (Gautier et al. 2013), we need to account for both sampling and read uncertainty arising from finite sample size and read counts, respectively, resulting in a mixture data model of two binomial distributions (Coop et al. 2010; Günther and Coop 2013) (Figs. S2 and S3). An advantage of the pool-seq method is that library preparation costs are much lower than those for individual-based sequencing methods. However, in pool-seq data, all individual-level information is lost, making it challenging to reuse the data for other analyses such as pedigree reconstruction.

Our process model can also be improved by accounting for additional evolutionary mechanisms that are currently missing. Our process model assumes that the allele frequencies in demes are independent. However, allele frequencies in demes may not be independent due to coupling via migration. This can yield false positives due to spurious correlations between allele frequencies and environmental gradients (Coop et al. 2010; Frichot et al. 2013). Another potential avenue for improvement is to build a process model that incorporates polygenic adaptation in which multiple loci regulate a phenotypic trait (such as body size). Lotterhos (2023) showed that because of genetic redundancy, a clinal pattern in a phenotypic trait may not always correspond to a clinal pattern in loci that regulate the trait. Consequently, our statistical model may fail to detect adaptive variation.

## Acknowledgements

We thank Hooten, Ruegg, and Zavaleta lab members for their feedback on the manuscript. Funding for this research was provided by NSF 2222525, NSF 1927177, NSF 2222524, and NSF 2222526. This work used Texas Advanced Computing Center (TACC) at The University of Texas at Austin through allocation TG-BIO250092 and TG-BIO260362 from the Advanced Cyberinfrastructure Coordination Ecosystem: Services & Support (ACCESS) program, which is supported by NSF 2138259, NSF 2138286, NSF 2138307, NSF 2137603, and NSF 2138296. We also thank Timothy H. Keitt for contributing computational resources for the project.

## Author Contributions

Nikunj Goel and Mevin B. Hooten conceived and designed the study with substantial feedback from Kristen C. Ruegg and Christen M. Bossu. Kristen C. Ruegg and Christen M. Bossu provided the genetic and climate data. Timothy M. Brown, Erika S. Zavaleta, Erica Robertson and Reza Goljani Amirkhiz collected Sierra Nevada Gray-crowned

Rosy-Finch samples. Ben J. Vernasco and Peri E. Bolton collected Wallowa samples in Oregon. Nikunj Goel conducted the mathematical and data analysis, Seorim Yi constructed the dyadic regression model of migration, and Nikunj Goel wrote the paper with feedback from Christen M. Bossu, Ben J. Vernasco, Erika Zavaleta, Kristen C. Ruegg, and Mevin B. Hooten. All authors reviewed and approved the manuscript.

