## Supplementary Information for "Identifying adaptive variation in spatially structured populations using low-coverage whole-genome sequencing data"

### Methods for data processing

#### *Sampling*

We compiled previously analyzed Rosy-Finch sequences and newly sampled, Sierra Gray-crowned and Wallowa Gray-crowned Rosy-Finch individuals. First, we included 113 individuals from 11 Brown-capped Rosy-Finch (BCRF) populations previously analyzed by DeSaix et al. (2022) and Funk et al. (2021), 3 individuals from 1 population of the Black Rosy-Finch (BLRF) (Funk et al. 2021), and 16 individuals from 5 Gray-crowned Rosy-Finch (GCRF) populations (Funk et al. 2021). Second, we expanded our sampling to include 52 Gray-crowned Rosy-Finch individuals from 6 additional populations in the California Sierras and 8 individuals from the Wallowa subspecies (WARF) in Oregon. The genetic data can be accessed using NCBI BioProj #: PRJNA1405131, PRJNA1297057, PRJNA659436, and PRJNA1380501.

#### *Sequencing*

DNA was extracted with the Qiagen DNeasy Blood and Tissue Kit following a modified protocol to maximize DNA yield from feathers (Schweizer and DeSaix 2023) and quantified with the Qubit dsDNA HS Assay Kit (Thermo Fisher Scientific, USA). We used a modified version of Illumina's Nextera Library Preparation protocol (*66*) to prepare whole-genome sequencing libraries and pooled the libraries by equal mass prior to sequencing. The pooled libraries were sequenced on two Illumina NovaSeq 6000 lanes at Novogene Corporation Inc.

#### *Read data processing*

To achieve reproducibility, we automated the analysis process with a Snakemake workflow management system (Mölder et al. 2025). We used *mega-non-model-wgs-snakeflow* (https://github.com/eriqande/mega-non-model-wgs-snakeflow) to process and map raw reads to the Brown-capped Rosy-finch genome assembly (Leucosticte australis, GCA_025504685.1), and to detect variants parallelizing across the genome using GATK v4.2.6.1 (McKenna et al. 2010; Van der Auwera et al. 2013). The *mega-non-model-wgs-snakeflow* workflow requires a "units" file, that links which samples were sequenced to the fastq files sequenced per sample, library and barcode information. The *mega-non-model-snakeflow* first trimmed pair-end fastq sequences using *fastp* (Chen et al. 2018), trimming Illumina adapter sequences and poly-G tails using a sliding window approach (SLIDINGWINDOW:4:20). We then used BWA 0.7.17 (Li and Durbin 2009) with the bwa-mem2 Snakemake wrapper (v1.23.3/bio/bwa/mem) to map trimmed sequences to the Brown-capped Rosy-finch genome (GCA_025504685.1). During mapping, we sorted and converted the resulting SAM files to BAM files using SAMtools 1.16 (Danecek et al. 2021; Li et al. 2009) and added read groups with Picard 3.0.0 (Broad Institute 2018). We then marked PCR duplicates with SAMtools (Danecek et al. 2021; Li et al. 2009) using the Snakemake wrapper v1.1.0/bio/picard/markduplicates. Finally, we used bamUtils with a poolSize of 10,000,000 and the parameter –poolSkipClip to clip overlaps in the BAM files. We calculated average sequencing depth using SAMtools stat (Li et al 2009, Danecek et al. 2021) to estimate the number of bases relative to the total genome length. To reduce sequence depth variation in rosy-finches, we followed the recommendations of (Lou and Therkildsen 2022) and downsampled the resulting BAM files to 5× coverage, using the subsample function in SAMtools (Li et al 2009, Danecek et al. 2021).

We generated gVCF files from downsampled BAM files using HaplotypeCaller in GATK 4.2.6.1 (McKenna et al. 2010; Van der Auwera et al. 2013). Given the early sequencing of rosy-finches was conducted on the Illumina HiSeq platform and current sequencing was completed on the Illumina NovoSeq 6000 platform, we included a minimum base quality score (*--min-base-quality-score*) of 33 and a minimum mapping quality score (*--minimum-mapping-quality-score*) of 20 for NovoSeq sequenced individuals, and minimum base quality score of 20 for HiSeq sequenced individuals. To accelerate computation, we generated genomic databases in ~3 Mb intervals across the genome and combined and indexed the genotyped VCF files with BCFtools 1.16 (Danecek et al. 2021). We used the *setGT* function in BCFtools to mark individuals with a read depth of 0 or PL of 0,0,0 as missing data. We applied a hard filter to the subsequent VCF file with the following parameters, "QD < 2.0 || FS > 60.0 || MQ < 40.0 || MQRankSum < -12.5 || ReadPosRankSum < -8.0", filtering the indels separately with "QD < 2.0 || FS > 200.0 || ReadPosRankSum < -20.0" to remove systematic errors as outlined in GATK best practices (Van der Auwera et al. 2013). We then used the *view* function in BCFtools to retain biallelic sites (-m 2 -M 2) that were missing in fewer than 20% of the sampled individuals ('F_MISSING < 0.20'), had a minor allele frequency of at least 0.05 (--min-af 0.05, --max-af 0.95), and had a sequencing quality score of at least 30 (‘QUAL > 30’) (*67*). To further remove potential platform batch effects, we conducted an $F_{ST}$-outlier scan in vcftools (Danecek et al. 2021) using individuals sequenced from the White Mountain Gray-crowned Rosy-Finch population on each sequencing platforms ($n_{\_NovoSeq}$=11 and $n_{\_HiSeq}$=8, including 4 duplicate individuals), and removed sites with a $F_{ST} > 0.05$.

#### *Ranking Environmental Variables with Gradient Forest*

To encompass the range of bioclimatic factors that may influence the Rosy-Finch alpine breeding habitat, we obtained 32 bioclimatic variables from the AdaptWest Project at a 1 km resolution (AdaptWest Project, 2022;Wang et al. 2016). To represent the current time period that corresponds to our sampled data, we obtained the bioclimatic variables as means across the time period of 1991–2020, and we obtained the dataset at an appropriate resolution for Rosy-Finch breeding movements (1 km). This 30-year time window is commonly used as a baseline in many studies because it allows us to average out idiosyncratic yearly fluctuations in the climate time series. However, this baseline may underestimate contemporary environmental gradients due to dramatic climate change over the preceding three decades.

In addition, we extracted snowpack data from MODIS/Terra Snow Cover Monthly L3 Global dataset (Hall and Riggs 2021) during the breeding season (May, June, and July) from 2020-2024. We accounted for potential variation in monthly snow cover by averaging snow cover for each month separately, as well as across the entire breeding season for each site. We also obtained data for elevation (SRTM; Farr et al. 2007) and soil water content (Trabucco and Zomer 2019), or the amount of water in soil, crucial for plants, hydrology, and climate.

We used the gradient forest algorithm to describe the associations of spatial environmental and genetic variables and identify the environmental predictors driving observed changes in allele frequency (Ellis et al. 2012; Fitzpatrick and Keller 2015). We fit gradient forest models to all the environmental variables listed above for the 24 sites with sample sizes of at least 3 individuals using the package gradientForest (Smith and Ellis 2013). In keeping with site grouping method that DeSaix et al. (2022) utilized with Brown-capped Rosy-Finch, separate sites were only combined into a single sampling unit based on their proximity and sample size. For example, individuals from the Lost Man Lake and Independence Lake sites were combined as a single sampling unit for subsequent analyses based on their proximity (<1 km) and the low sample sizes (5 and 1 individuals, respectively; Supplemental Table 1). Due to the large SNP dataset, we ran each model 5 times with a randomly sampled subset of 50,000 SNPs, as the response variables. We used the weighted $r^2$ value from the output for each model to assess the importance of the predictor variables. The top uncorrelated variables with correlation less than equal to 0.7 were selected for use in our subsequent hierarchical Bayesian genotype-environmental association analysis.

*Identifying adaptive genes*

To identify genes associated with significant SNPs from our model, we utilized BEDTools (Quinlan and Hall 2010) *closest* function with a previously created Brown-capped Rosy Finch genome annotation (Funk et al. 2023) to identify genes close to the candidate SNP positions. The output from this tool was filtered to identify only protein-coding genes within 25kb of a given SNP. For plotting purposes, we used nucmer (Marçais et al. 2018) to align our Brown-capped Rosy-Finch genome assembly scaffolds into pseudochromosomes, based on synteny with the Zebra Finch (*Taeniopygia guttata*) genome assembly (GenBank: GCF_003957565.2). We then used the resulting chromosome positions when plotting SNPs in the Manhattan plots.

### Analysis of RAD-seq data

To analyze RAD-seq data, we modified the data model in the main text to estimate allele frequencies and coalescent times using called-genotypes at $L$ polymorphic loci. We assume that the genotype of the $i$th individual in sampling site $k$ and loci $l$ is $g_{ilk}$. We split the genetic dataset into two parts—$L_1$ loci to model allele frequencies and $L_2$ loci to model coalescent times. We define an allele count variable for $L_1$ loci as

$$\tilde{y}_{lk} = \sum_{i=1}^{\tilde{N}_{lk}} g_{ilk}.$$

The variable $\tilde{y}_{lk}$ corresponds to the total number of reference alleles in the sample for $\tilde{N}_{lk}$ individuals that were genotyped at loci $l$. Note that in the data model for genotype-likelihoods, total number of reference alleles was treated as parameter as the genotype of an individual is unknown. To account for missing genotypes, we assumed that $\tilde{N}_{lk}$ is less than the total number of individuals that were sampled at site $k$ ($N_k$). Assuming the individuals were sampled randomly, we can statistically model the reference allele frequencies at $L_1$ loci using a binomial distribution

$$\tilde{y}_{lk} \sim \text{Binomial}\left(2\tilde{N}_{lk}, p_{lk}\right).$$

Next, we integrated over allele frequencies in equation (20) to obtain a joint data and process model:

$$\tilde{y}_{lk} \sim \frac{{}_1F_1\left(\mu_{lk}^{(2)}\kappa_{lk}^{*} + \tilde{y}_{lk}, 2\tilde{N}_{lk} + \kappa_{lk}^{*}, \tilde{s}_{lk}\right)}{{}_1F_1\left(\mu_{lk}^{(2)}\kappa_{lk}^{*}, \kappa_{lk}^{*}, \tilde{s}_{lk}\right)} \text{BetaBinomial}\left(\tilde{y}_{lk} | 2\tilde{N}_{lk}, \mu_{lk}^{(2)}, \kappa_{lk}^{*}\right),$$

To estimate coalescent times, we calculate genetic dissimilarity ($D_{kk'}$) using $L_2$ loci, which can then be used to estimate migration rates using equations (9), (21), and (22) in the main text. Finally, we tested the RAD-seq data model using synthetic genotypes generated in step three (see Model Testing section) corresponding to the data used to generate Fig. 4A (see Fig. S1).

### Analysis of pool-seq data

To analyze pool-seq data, we modified the data model in the main text to estimate allele frequencies and coalescent times using pool-seq data. We assumed that the DNA is pooled from $N_k$ individuals at sampling site $k$. These pooled samples are sequenced at $L$ polymorphic loci, yielding a total number of $R_{lk}$ reads at locus $l$ with $r_{lk}$ reads corresponding to the reference allele. To model allele frequencies and coalescent times, we split the genetic dataset into two parts: $L_1$ loci to model allele frequencies and $L_2$ loci to model coalescent times. Assuming that individuals were sampled randomly, we statistically modeled reference allele frequencies at $L_1$ loci as a mixture of two binomial distributions (Günther and Coop 2013):

$$r_{lk} \sim \sum_{n=0}^{2N_k} \text{Binomial}\left(r_{lk} \mid R_{lk}, \frac{n}{2N_k}\right) \text{Binomial}(n \mid 2N_k, p_{lk}).$$

The first binomial distribution corresponds to the model of read data, and the second binomial distribution corresponds to the model of sampling individuals to form a sequencing pool.

There are two possible options to integrate the above data model with the process model in equation (20). Option one is to precompute $\text{Binomial}\left(r_{lk} \mid R_{lk}, \frac{n}{2N_k}\right)$ and treat it as $a_{nlk}$ in equation (2). The advantage of this option is that we can follow the calculations in the main text to estimate effective sample size and effective counts of reference alleles (Eqs. 4-8; see Fig. S3), which can then be subsequently used as data in equation (29). This option effectively allows us to use the same pipeline as genotype-likelihoods. The second option is to marginalize over only $p_{lk}$ to derive the following joint data and process model:

$$r_{lk} \sim \sum_{n=0}^{2N_k} \frac{{}_1F_1\left(\mu_{lk}^{*}\mu_{lk}^{(2)} + n, 2N_k + \kappa_{lk}^{*}, \tilde{s}_{lk}\right)}{{}_1F_1\left(\mu_{lk}^{(2)}\kappa_{lk}^{*}, \kappa_{lk}^{*}, \tilde{s}_{lk}\right)} \text{Binomial}\left(r_{lk} \mid R_{lk}, \frac{n}{2N_k}\right) \text{BetaBinomial}\left(n \mid 2N_k, \mu_{lk}^{(2)}, \kappa_{lk}^{*}\right).$$

This joint likelihood model is more accurate than option one but can be very slow due to the summation. Finally, to estimate mean migration rates, we use $\hat{p}_{lk} = r_{lk}/R_{lk}$ to calculate genetic dissimilarity ($D_{kk'}$), which is then linked to coalescent times using equation (9).

To test the pool-seq data model, we generated and analyzed synthetic read data using allele counts obtained from step three of the Model Testing section, corresponding to the data in Fig. 4A (see Fig. S2). To generate read data, we sampled from $\text{Binomial}\left(R_{lk}, \frac{n_{lk}}{2N_k}\right)$, where $R_{lk}$ is a Poisson random variable with mean equal to sequencing-depth (=4) times the number of individuals sampled (=10), and $n_{lk}$ is the total number of reference alleles for locus $l$ in the deme $k$. We used effective sample size and effective counts of reference alleles to fit the model because it was much faster than option two.

### Covariates for dyadic regression

To constrain migration rates, we specified a dyadic regression model (Schwob et al. 2024)

$$M_{kk'} \sim \text{Exponential}\left(e^{\gamma_0 + \boldsymbol{x}_{kk'}^T \boldsymbol{\gamma}}\right).$$

We let $\boldsymbol{\gamma}$ represent the regression coefficients and $\boldsymbol{x}_{kk'}$ represent the dyadic covariates integrated over the least cost path from deme $k$ to deme $k'$. To obtain the line integrated covariates ($\boldsymbol{x}_{kk'}$), we summed the environmental variables (mean temperature difference between warmest and coldest month, mean summer precipitation, precipitation as snow, Hargreave's climatic moisture index, mean annual relative humidity, and elevation) along shortest path on land (at 0.05 degree resolution) and divide it by distance along the shortest path. These covariates are then standardized to have a zero mean and a standard deviation of one. In addition, we also considered the shortest distance (standardized) and its interaction with other environmental variables as additional covariates.

### Full Bayesian likelihood

In the main text, we used effective sample sizes and effective counts of reference alleles as data to fit the structured genotype-likelihood method (Eq. 29). However, the 'true' joint data and process likelihood is given by

$$\boldsymbol{y}_{lk} \sim \sum_{n=0}^{2N_{lk}} a_{nlk} \frac{{}_1F_1\left(\mu_{lk}^{(2)}\kappa_{lk}^{*} + n, 2N_{lk} + \kappa_{lk}^{*}, \tilde{s}_{lk}\right)}{{}_1F_1\left(\mu_{lk}^{(2)}\kappa_{lk}^{*}, \kappa_{lk}^{*}, \tilde{s}_{lk}\right)} \text{BetaBinomial}\left(n|2N_{lk}, \mu_{lk}^{(2)}, \kappa_{lk}^{*}\right).$$

To test this likelihood, we re-analyzed the data in Figure 4A. The results (Fig. S7) are qualitatively consistent with the results in Figure 4A. However, our simulations required 3.5 times more time to fit the 'true' likelihood.

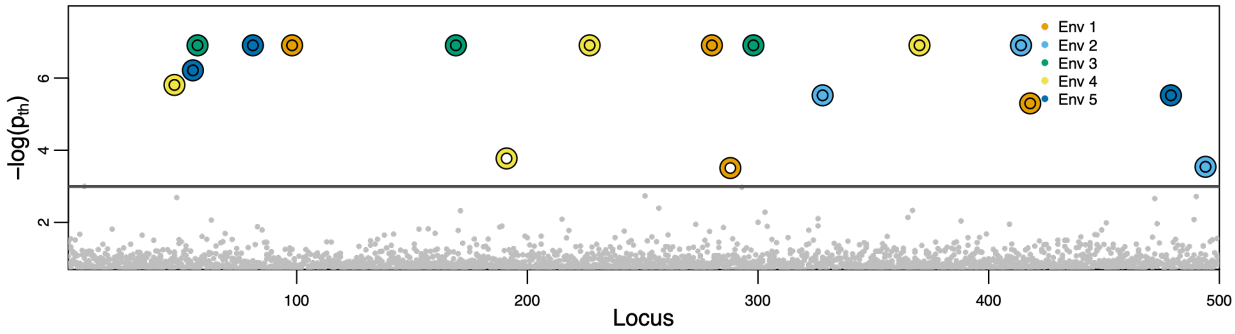


**Figure S1:** Manhattan plot showing negative log probability $(-\log(p_{\mathrm{th}}))$that the posterior distribution of sensitivity coefficients $(\beta_{lj})$ includes zero for RAD-seq synthetic data. The points above the black horizontal line have $p_{\mathrm{th}}$ less than 0.05. We denote points above the black line using an outer ring and a center. The color of the center (ring) corresponds to the true (statistically inferred) environmental variable responsible for local adaptation. The underlying synthetic data for generating this plot is same as Fig. 4A.

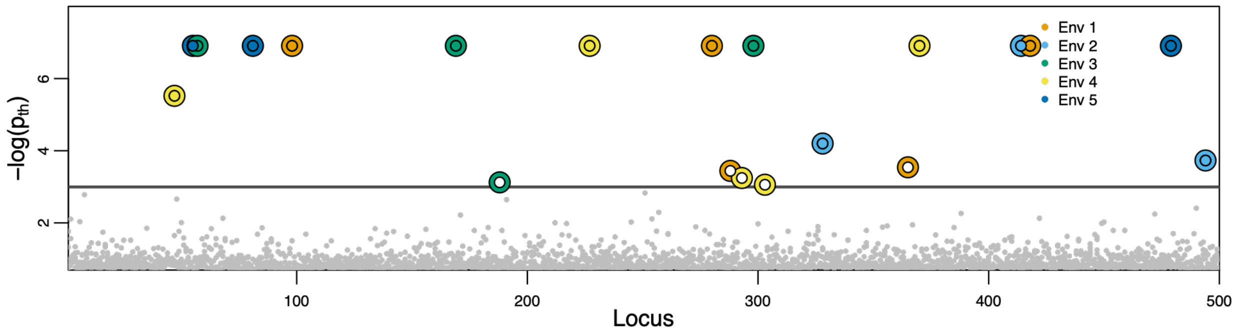


**Figure S2:** Manhattan plot showing negative log probability $(-\log(p_{\text{th}}))$ that the posterior distribution of sensitivity coefficients $(\beta_{lj})$ includes zero for pool-seq synthetic data. The points above the black horizontal line have $p_{\text{th}}$ less than 0.05. We denote points above the black line using an outer ring and a center. The color of the center (ring) corresponds to the true (statistically inferred) environmental variable responsible for local adaptation. The underlying synthetic data for generating this plot is same as Fig. 4A.

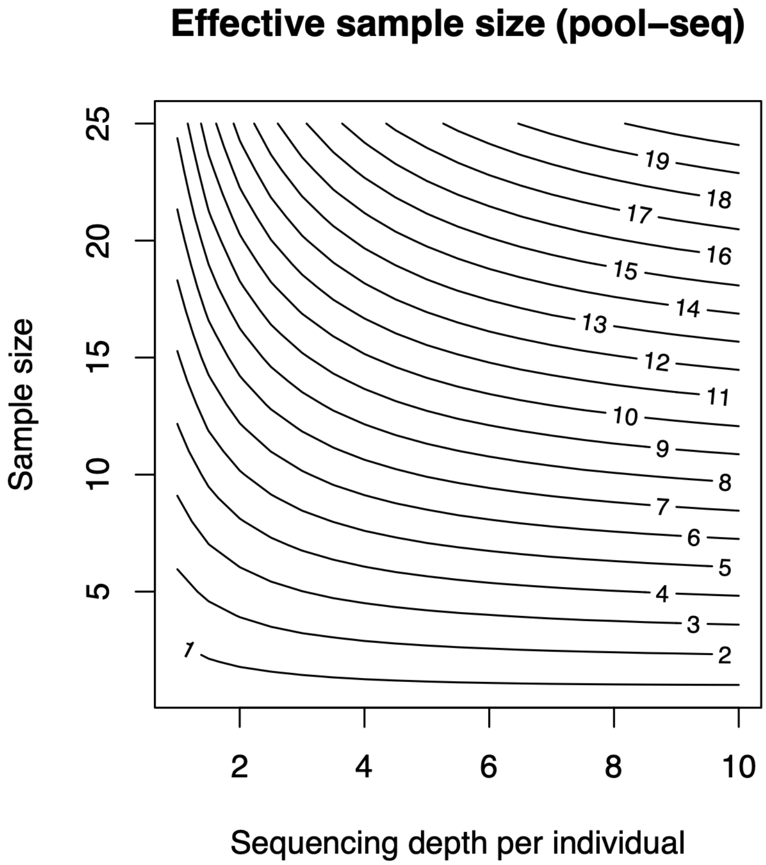


**Figure S3:** Effective sample size as a function of sequencing depth per individual and number of individuals sampled at a location to form the pool. The contours represent combinations of sequencing depth and sample size for which the effective sample size is constant. Based on the desired level of precision in estimates of allele frequency (determined by the effective sample size), a biologist can make an informed decision about the relative allocation of resources between collecting individuals and sequencing effort. Note that for the same sequencing effort, the effective sample size for low-coverage whole-genome sequencing data (Fig. 3 in the main text) is higher than that for pool sequencing data.

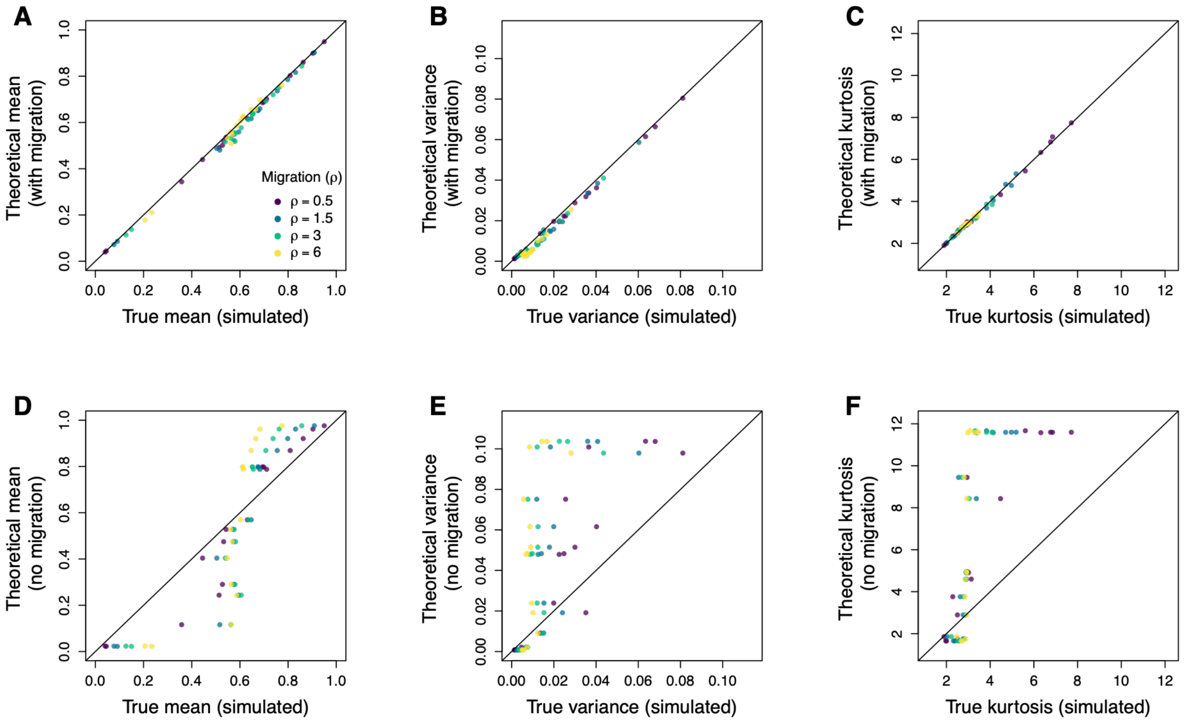


**Figure S4:** Comparing the means (**A** and **D**), variances (**B** and **E**), and kurtoses (**C** and **F**) of analytically derived and simulated stationary distributions of allele frequencies. Plots (**A**), (**B**) and (**C**) shows that equation (16) reasonably approximates the 'true' stationary distribution, which is obtained by simulating stochastic evolutionary dynamics in equation (11). In contrast, plots (**D**), (**E**) and (**F**) shows that the analytically derived stationary distribution for the 'no migration' case (Goel et al. 2025) deviates substantially from the 'true' stationary distribution. These plots suggest that migration-selection balance can influence spatial genetic variation. Thus, statistical methods that account for migration may yield better inferences. To generate these plots, we use migration rates inferred from Rosy Finch data (Fig. 5). We used $\mu_l = 0.5$ , $\kappa_l = 1.4$, and $\beta_{lj} = 15$ for all plots, where $j$ corresponds to the mean summer precipitation.

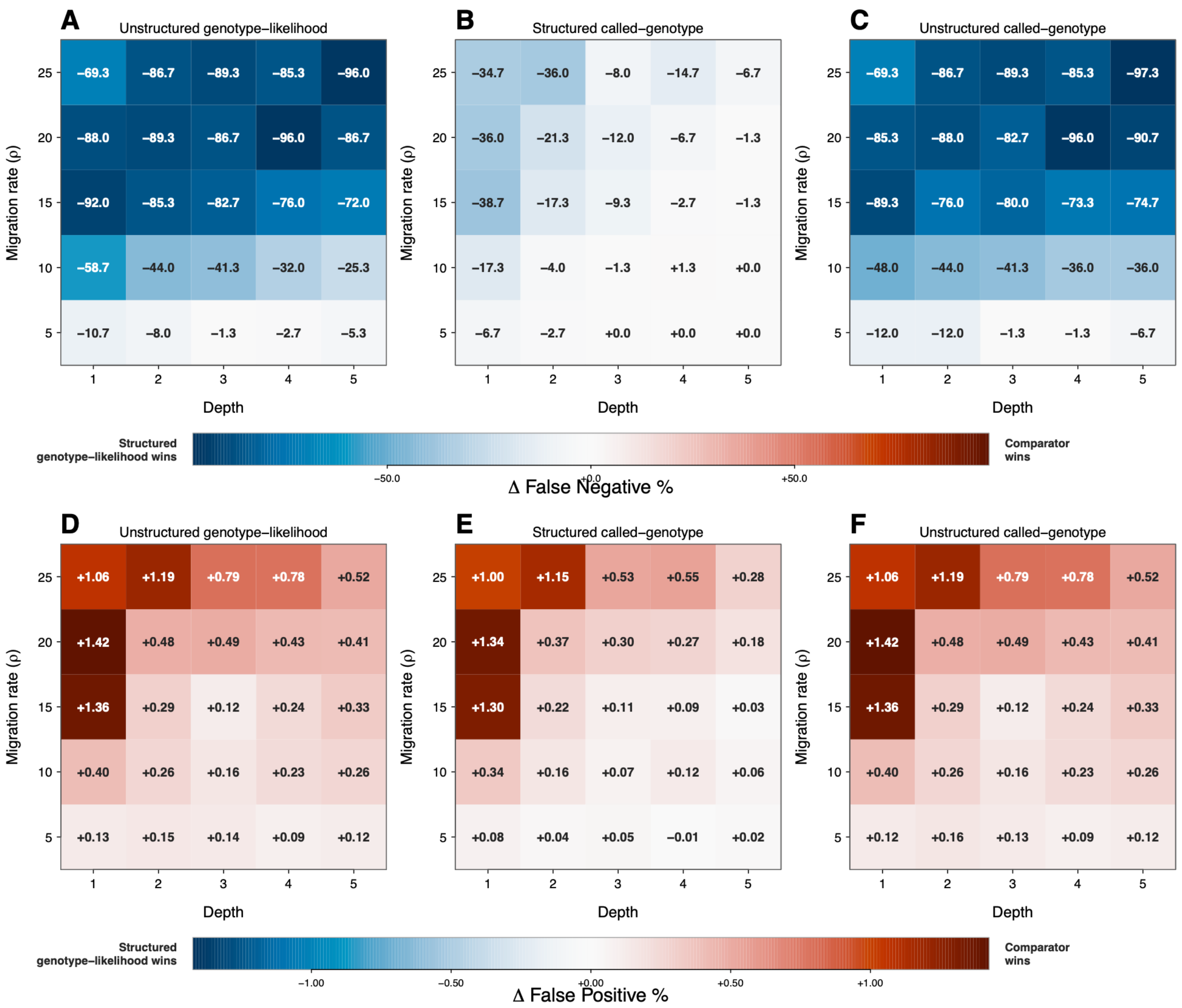


**Figure S5:** Difference in false-negative (**A**, **B**, and **C**) and false-positive (**D**, **E**, and **F**) percentages between the base model (structured genotype-likelihood) and the unstructured genotype-likelihood model (**A** and **D**), between the base model (structured genotype-likelihood) and the structured called-genotype model (**B** and **E**), and between the base model (structured genotype-likelihood) and the unstructured called-genotype model (**C** and **F**). As expected, the structured genotype-likelihood model performed best in high migration and low sequencing depth regimes, underscoring the importance of accounting for genotype uncertainty and structured migration for identifying adaptive variation. We define false negative % as false negatives/adaptive loci x 100 and false positive % as false positives/neutral loci x 100.

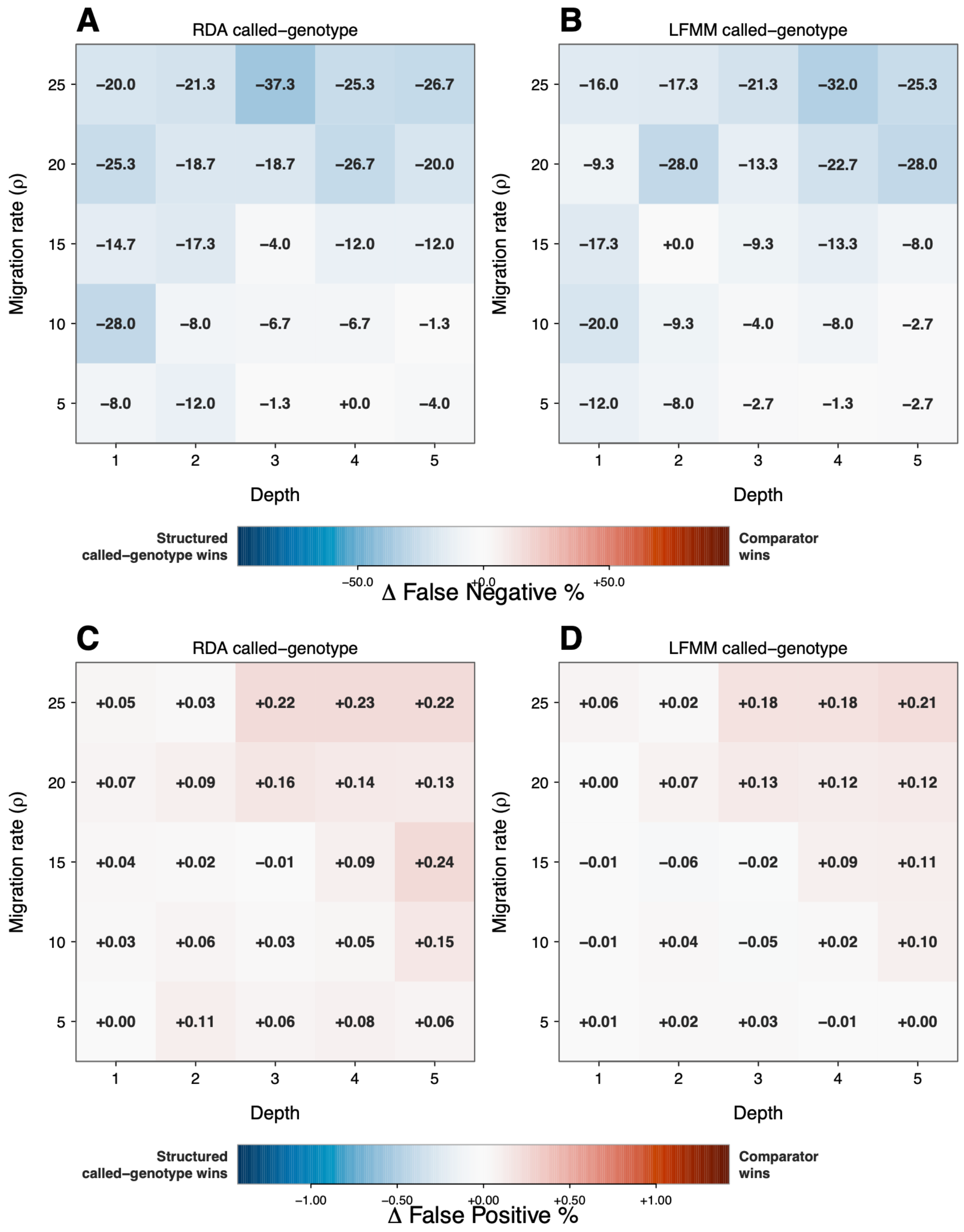


**Figure S6:** Difference in false-negative (**A** and **B**) and false-positive (**C** and **D**) percentages between the base model (structured called-genotype) and the RDA called-genotype model (**A** and **C**), and between the base model (structured called-genotype) and the LFMM called-genotype model (**B** and **D**). As expected, the structured called-genotype model performed best in high migration and low sequencing depth regimes, underscoring the importance of accounting for genotype uncertainty and structured migration when identifying adaptive variation. We define false negative % as false negatives/adaptive loci x 100 and false positive % as false positives/neutral loci x 100.

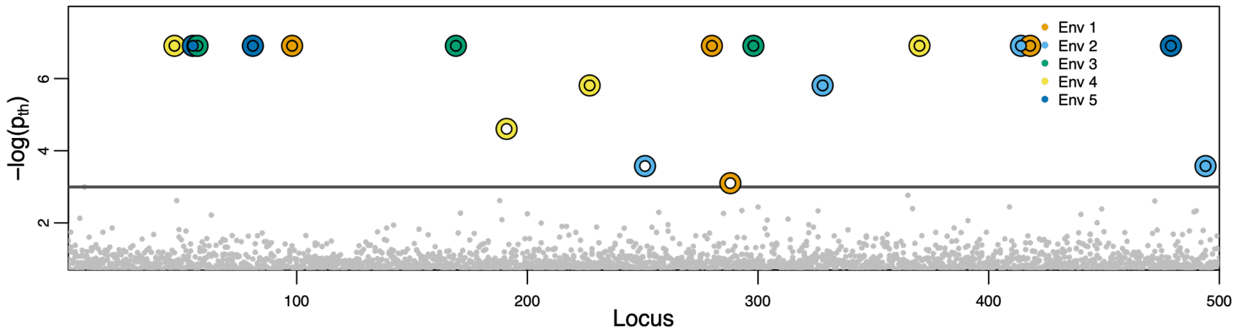


**Figure S7:** Manhattan plot showing negative log probability ($-\log(p_{\mathrm{th}})$) that the posterior distribution of sensitivity coefficients ($\beta_{lj}$) includes zero for synthetic data. The data was analyzed using the full Bayesian model in which migration rates were treated as parameters rather than known inputs. The points above the black horizontal line have $p_{\mathrm{th}}$ less than 0.05. We denote points above the black line using an outer ring and a center. The color of the center (ring) corresponds to the true (statistically inferred) environmental variable responsible for local adaptation. The underlying synthetic data for generating this plot is same as Fig. 4A.

**Tables**

| Species | State | Location | Climate Group | Lat | Lon | $N_k$ | Reference |
|---|---|---|---|---|---|---|---|
| BLRF | MT | Beartooth Plateau | Beartooth Plateau | 44.99 | -109.57 | 3 | Funk et al. (2021) |
| GCRF | AK | Dalton Highway | Dalton Highway | 68.20 | -149.41 | 3 | Funk et al. (2021) |
| BCRF | CO | Devil's Causeway | Devils Causeway | 40.03 | -107.17 | 14 | DeSaix et al. (2022) |
| BCRF | CO | Emma Burr Mountain | Emma Burr Mountain | 38.75 | -106.42 | 3 | DeSaix et al. (2022) |
| BCRF | CO | Engineer Mountain | Engineer Mountain | 37.97 | -107.58 | 8 | DeSaix et al. (2022) |
| BCRF | CO | Horseshoe Basin | Engineer Mountain | 37.95 | -107.55 | 18 | DeSaix et al. (2022) |
| GCRF | CA | Humphrey Basin | Humphrey Basin | 37.23 | -118.68 | 14 | Current Sampling |
| BCRF | CO | Independence Lake* | Emma Burr Mountain | 39.14 | -106.57 | 1 | DeSaix et al. (2022) |
| BCRF | CO | Lake Agnes | Lake Agnes | 40.47 | -105.90 | 15 | DeSaix et al. (2022) |
| BCRF | CO | Lost Man Lake* | Emma Burr Mountain | 39.15 | -106.57 | 5 | DeSaix et al. (2022) |
| BCRF | WY | Medicine Bow NF | Medicine Bow NF | 41.37 | -106.30 | 12 | DeSaix et al. (2022) |
| BCRF | CO | Mountain Maxwell | Mountain Maxwell | 37.25 | -105.15 | 7 | DeSaix et al. (2022) |
| BCRF | CO | Mt. Evans | Arapaho and Roosevelt NF | 39.59 | -105.64 | 10 | DeSaix et al. (2022) |
| GCRF | AK | Mt. Prindle | Mt. Prindle | 65.44 | -146.54 | 4 | Funk et al. 2021 |
| BCRF | CO | Pike's Peak | Pike's Peak | 38.83 | -105.04 | 20 | DeSaix et al. (2022) |

| | | | | | | | |
|---|---|---|---|---|---|---|---|
| GCRF | CA | Piute Pass | Humphrey Basin | 37.24 | -118.47 | 8 | Current Sampling |
| GCRF | CA | Sierra Nevada | Humphrey Basin | 36.77 | -118.39 | 1 | Current Sampling |
| GCRF | AK | Sugarloaf Mt. | Sugarloaf Mt. | 63.81 | -148.83 | 1 | Funk et al. (2021) |
| GCRF | CA | Sweetwaters Mountains | Sweetwaters Mountains | 38.42 | -119.29 | 5 | Current Sampling |
| GCRF | AK | Valdez-Cordova | Valdez-Cordova | 63.15 | -146.28 | 4 | Funk et al. (2021) |
| GCRF | CA | Virginia Lakes | Sweetwaters Mountains | 38.05 | -119.26 | 11 | Current Sampling |
| WARF | OR | Wallowa | Wallowa | 45.57 | -117.53 | 8 | Current Sampling |
| GCRF | CA | White Mountain | Humphrey Basin | 37.63 | -118.25 | 13 | Current Sampling |
| GCRF | CA | Yosemite NP | Sweetwaters Mountains | 37.92 | -119.24 | 4 | Funk et al. (2021) |

**Table S1:** List of sampled species, their geographical locations (state, local landmark, and coordinates), climatic grouping for the hierarchical Bayesian model herein, number of sampled individuals, and citations for corresponding references.

*Locations combined for gradient forest analysis to identify uncorrelated environmental variables

| | SWC | MSP | RH | Cn | MJS |
|---|---|---|---|---|---|
| **SWC** | — | +0.082 | -0.576 | -0.490 | -0.077 |
| **MSP** | +0.082 | — | -0.171 | +0.344 | -0.700 |
| **RH** | -0.576 | -0.171 | — | +0.433 | +0.233 |
| **TD** | -0.490 | +0.344 | +0.433 | — | -0.453 |
| **July snow** | -0.077 | -0.700 | +0.233 | -0.453 | — |

**Table S2:** Pairwise correlation between environmental covariates. SWC is soil water content, MSP is mean summer precipitation (mm), RH is relative humidity, Cn is continentality, and MJS is mean July snowpack.

| Gene | Function | Reference |
|---|---|---|
| **Pigmentation** | | |
| **AP3B1** | Melanosome biogenesis, TYR recruitment in rosy finches, and melanosome organization (GO term); Differential expression between Swainson's thrush in the migratory and non-migratory phase of the annual cycle. Differentially expressed between willow warblers with different migratory behavior. | Boss et al. (2016); Funk et al. (2023); and Johnston et al. (2016) |
| **APC** | Associated with plumage color variation in yellow and red-shafted flickers. | Aguillon et al. (2021) |
| **ASIP** | Associated with plumage color variation in passerines and with melanin biosynthetic processes (GO term) | Funk and Taylor (2019); Nadeau et al. (2008); and Ruegg et al. (2014) |
| **EDN3** | Melanocyte production and involved in melanocyte differentiation (GO term) | Funk et al. (2023); and Saldana-Caboverde and Kos (2010) |
| **LRMDA** | Involved in melanocyte differentiation (GO term) and plays a role in eumelanin synthesis; Mutation that results in albinism in humans | Grønskov et al. (2013) |
| ***NDUFAF2*** | Crown coloration | Funk et al. (2023) |
| **TYRP1** | Associated with the melanosome (GO term) | Nadeau et al. (2007); and Skoglund and Höglund (2010) |
| **Migration** | | |
| **ATG2B** | Genetic variants underlying variation in migration in blackcaps | Delmore et al. (2020) |
| **FRMD3** | Differential expression between Swainson's thrush in the migratory and non-migratory phase of the annual cycle, and higher expression in fat line of broiler chickens | Johnston et al. (2016); and Wang et al. (2021) |

| | | |
|---|---|---|
| **ICA1** | Differential expression between Swainson's thrush in the migratory and non-migratory phase of the annual cycle; eggshell color in broiler chickens | Johnston et al. (2016); and Yang et al. (2023) |
| **KLF3** | Regulation of adipogenesis and lipid metabolism, affects chicken abdominal fat content | Raza et al. (2022); and Wang et al. (2020) |
| **PRKAG2** | Differentially expressed in Swainson's thrush with different migratory states in the Cluster N brain region, a visual area of the brain that is activated during migration | Louder et al. (2024) |
| **Physiology linked to high altitude adaptation** | | |
| **AGGF1** | High altitude-adaption gene under selection in Tibetan and Dahe pigs; Angiogenic factor promoting proliferation of endothelial cells | Dong et al. (2014); and Funk et al. (2023) |
| **ALDH1A1** | Retinol metabolism and metabolic responses to high-fat diet | Funk et al. (2023); and Kiefer et al. (2012); |
| **EGLN1** | High altitude adaptation that is a key regulator in the body's response to oxygen levels by controlling the breakdown of a protein called hypoxia-inducible factor (HIF) | Aggarwal et al. (2015); and Funk et al. (2023) |
| **EPAS1** | Key gene mutated in Tibetan populations adapted to living at high altitudes, and convergent evolution of EPAS1 identified in high altitude ducks | Beall et al. (2010); Graham and McCracken (2019); Simonson et al. (2010); and Yi et al. (2010) |
| **JMY** | Stress responsive protein upregulated during hypoxia | Coutts et al. (2011); and Funk et al. (2023); |
| **PPARGC1A** | Associated with cold tolerance, heat stress, and water scarcity (alpine adaptation) in chickens; PGC-1α is highly expressed in tissues with active oxidative metabolism, such as brown adipose tissue (BAT), heart, and skeletal muscle. Differentially expressed in fasting goose liver, in relation to fat metabolism | Chen et al. (2021); Gheyas et al. (2021); and Rius-Pérez et al. (2020) |
| **RTEL1** | Association analyses in chickens identified RTEL1 as associated with body temperature changes, suggesting tolerance when subjected to acute heat stress; RTEL1 | Rong et al. (2017); and Zhuang et al. (2020) |

| | | |
|---|---|---|
| | variants associated with a decreased risk of HAPE, high altitude pulmonary edema, in the Chinese Han population | |
| **TNFSF11** | TNFSF11 encodes the RANKL protein, which regulates bone remodeling, osteoclast differentiation, and immune responses. In hypoxic environments HIF1α serves as a promoter in RANKL-induced osteoclast differentiation | Wang et al. (2022) |
| **Physiology linked to stress response** | | |
| **COL22A1** | Significant expression related to heat stress in turkey | Reed et al. (2021) |
| **CRHBP** | CRHBP as the negative feedback regulator in the HPA axis (stress response) | Wan et al. (2022) |
| **GPX8** | Protecting cells from oxidative damage by neutralizing harmful reactive oxygen species (ROS). In chickens, GPX8 is upregulated in response to oxidative stress | Aryal et al. (2025); and Pei et al. (2023) |
| **Beak and feather development** | | |
| **FOXO3** | Developmental process of the skin and feather follicles in Zhedong white goose | Mabrouk et al. (2022) |
| **FZD4** | FZD4 is expressed in the growth and development of feather follicles for Anser anser and Anser cygnoides | Msuthwana et al. (2022) |
| **GLI3** | A key control gene within the Sonic hedgehog pathway that dictates whether an embryonic region develops a feather, hair follicle, scale. Differentially expressed in scale-to-feather conversion induced in chickens | Cooper and Milinkovitch (2023) |
| **LRRIQ1** | Associated with beak shape and size variation in Darwin's finches and great tits | Bosse et al. (2017) |
| **NELL1** | Associated with cold tolerance and muscle development and growth in chickens; Linked to bill morphology in great tits | Bosse et al. (2017); Fedorova et al. (2022); and Yin et al. (2019) |

**Table S3.** Candidate genes that were located within 25 kb of the statistically significant SNPs identified in our mechanistic model grouped into 5 broad functional categories (gray).

Aggarwal, S., A. Gheware, A. Agrawal, S. Ghosh, I. G. V. Consortium, B. Prasher, and M. Mukerji. 2015. Combined genetic effects of EGLN1 and VWF modulate thrombotic outcome in hypoxia revealed by Ayurgenomics approach. Journal of Translational Medicine 13:184.

Aguillon, S. M., J. Walsh, and I. J. Lovette. 2021. Extensive hybridization reveals multiple coloration genes underlying a complex plumage phenotype. Proceedings of the Royal Society B: Biological Sciences 288:20201805.

Aryal, B., J. Kwakye, O. W. Ariyo, A. F. Ghareeb, M. C. Milfort, A. L. Fuller, S. Khatiwada et al. 2025. Major oxidative and antioxidant mechanisms during heat stress-induced oxidative stress in chickens. Antioxidants 14:471.

Beall, C. M., G. L. Cavalleri, L. Deng, R. C. Elston, Y. Gao, J. Knight, C. Li et al. 2010. Natural selection on EPAS1 (HIF2α) associated with low hemoglobin concentration in Tibetan highlanders. Proceedings of the National Academy of Sciences 107:11459-11464.

Boss, J., M. Liedvogel, M. Lundberg, P. Olsson, N. Reischke, S. Naurin, S. Åkesson et al. 2016. Gene expression in the brain of a migratory songbird during breeding and migration. Movement Ecology 4:4.

Bosse, M., L. G. Spurgin, V. N. Laine, E. F. Cole, J. A. Firth, P. Gienapp, A. G. Gosler et al. 2017. Recent natural selection causes adaptive evolution of an avian polygenic trait. Science 358:365-368.

Chen, S., Y. Zhou, Y. Chen, and J. Gu. 2018. fastp: An ultra-fast all-in-one FASTQ preprocessor. Bioinformatics 34:i884-i890.

Chen, Z., Y. Xing, X. Fan, T. Liu, M. Zhao, L. Liu, X. Hu et al. 2021. Fasting and Refeeding Affect the Goose Liver Transcriptome Mainly Through the PPAR Signaling Pathway. The Journal of Poultry Science 58:245-257.

Cooper, R. L., and M. C. Milinkovitch. 2023. Transient agonism of the sonic hedgehog pathway triggers a permanent transition of skin appendage fate in the chicken embryo. Science Advances 9:eadg9619.

Coutts, A. S., I. M. Pires, L. Weston, F. M. Buffa, M. Milani, J. L. Li, A. L. Harris et al. 2011. Hypoxia-driven cell motility reflects the interplay between JMY and HIF-1α. Oncogene 30:4835-4842.

Danecek, P., J. K. Bonfield, J. Liddle, J. Marshall, V. Ohan, M. O. Pollard, A. Whitwham et al. 2021. Twelve years of SAMtools and BCFtools. Gigascience 10:giab008.

Delmore, K., J. C. Illera, J. Pérez-Tris, G. Segelbacher, J. S. Lugo Ramos, G. Durieux, J. Ishigohoka et al. 2020. The evolutionary history and genomics of European blackcap migration. eLife 9:e54462.

DeSaix, M. G., T. L. George, A. E. Seglund, G. M. Spellman, E. S. Zavaleta, and K. C. Ruegg. 2022. Forecasting climate change response in an alpine specialist songbird reveals the importance of considering novel climate. Diversity and Distributions 28:2239-2254.

Dong, K., N. Yao, Y. Pu, X. He, Q. Zhao, Y. Luan, W. Guan et al. 2014. Genomic scan reveals loci under altitude adaptation in Tibetan and Dahe pigs. PLoS One 9:e110520.

Ellis, N., S. J. Smith, and C. R. Pitcher. 2012. Gradient forests: Calculating importance gradients on physical predictors. Ecology 93:156-168.

Farr, T. G., P. A. Rosen, E. Caro, R. Crippen, R. Duren, S. Hensley, M. Kobrick et al. 2007. The shuttle radar topography mission. Reviews of Geophysics 45:RG2004.

Fedorova, E. S., N. V. Dementieva, Y. S. Shcherbakov, and O. I. Stanishevskaya. 2022. Identification of Key Candidate Genes in Runs of Homozygosity of the Genome of Two Chicken Breeds, Associated with Cold Adaptation. Biology 11:547.

Fitzpatrick, M. C., and S. R. Keller. 2015. Ecological genomics meets community-level modelling of biodiversity: Mapping the genomic landscape of current and future environmental adaptation. Ecology Letters 18:1-16.

Funk, E. R., N. A. Mason, S. Pálsson, T. Albrecht, J. A. Johnson, and S. A. Taylor. 2021. A supergene underlies linked variation in color and morphology in a Holarctic songbird. Nature Communications 12:6833.

Funk, E. R., G. M. Spellman, K. Winker, J. J. Withrow, K. C. Ruegg, and S. A. Taylor. 2023. The genetic basis of plumage coloration and elevation adaptation in a clade of recently diverged alpine and arctic songbirds. Evolution 77:705-717.

Funk, E. R., and S. A. Taylor. 2019. High-throughput sequencing is revealing genetic associations with avian plumage color. The Auk 136:ukz048.

Gheyas, A. A., A. Vallejo-Trujillo, A. Kebede, M. Lozano-Jaramillo, T. Dessie, J. Smith, and O. Hanotte. 2021. Integrated Environmental and Genomic Analysis Reveals the Drivers of Local Adaptation in African Indigenous Chickens. Molecular Biology and Evolution 38:4268-4285.

Goel, N., C. M. Bossu, J. J. Van Ee, E. Zavaleta, K. C. Ruegg, and M. B. Hooten. 2025. Identifying genomic adaptation to local climate using a mechanistic evolutionary model. Methods in Ecology and Evolution 16:2448-2460.

Graham, A. M., and K. G. McCracken. 2019. Convergent evolution on the hypoxia-inducible factor (HIF) pathway genes EGLN1 and EPAS1 in high-altitude ducks. Heredity 122:819-832.

Grønskov, K., Christopher M. Dooley, E. Østergaard, Robert N. Kelsh, L. Hansen, Mitchell P. Levesque, K. Vilhelmsen et al. 2013. Mutations in C10orf11, a Melanocyte-Differentiation

Gene, Cause Autosomal-Recessive Albinism. The American Journal of Human Genetics 92:415-421.

Günther, T., and G. Coop. 2013. Robust identification of local adaptation from allele frequencies. Genetics 195:205-220.

Hall, D., and G. Riggs. 2021. MODIS/Terra Snow Cover Monthly L3 Global 0.05Deg CMG, Version 61, NASA National Snow and Ice Data Center Distributed Active Archive Center.

Johnston, R. A., K. L. Paxton, F. R. Moore, R. K. Wayne, and T. B. Smith. 2016. Seasonal gene expression in a migratory songbird. Molecular Ecology 25:5680-5691.

Kiefer, F. W., G. Orasanu, S. Nallamshetty, J. D. Brown, H. Wang, P. Luger, N. R. Qi et al. 2012. Retinaldehyde Dehydrogenase 1 Coordinates Hepatic Gluconeogenesis and Lipid Metabolism. Endocrinology 153:3089-3099.

Li, H., and R. Durbin. 2009. Fast and accurate short read alignment with Burrows–Wheeler transform. Bioinformatics 25:1754-1760.

Li, H., B. Handsaker, A. Wysoker, T. Fennell, J. Ruan, N. Homer, G. Marth et al. 2009. The sequence alignment/map format and SAMtools. Bioinformatics 25:2078-2079.

Lou, R. N., and N. O. Therkildsen. 2022. Batch effects in population genomic studies with low-coverage whole genome sequencing data: Causes, detection and mitigation. Molecular Ecology Resources 22:1678-1692.

Louder, M. I. M., H. Justen, A. A. Kimmitt, K. S. Lawley, L. M. Turner, J. D. Dickman, and K. E. Delmore. 2024. Gene regulation and speciation in a migratory divide between songbirds. Nature Communications 15:98.

Mabrouk, I., Y. Zhou, S. Wang, Y. Song, X. Fu, X. Xu, T. Liu et al. 2022. Transcriptional Characteristics Showed That miR-144-y/FOXO3 Participates in Embryonic Skin and Feather Follicle Development in Zhedong White Goose. Animals 12:2099.

Marçais, G., A. L. Delcher, A. M. Phillippy, R. Coston, S. L. Salzberg, and A. Zimin. 2018. MUMmer4: A fast and versatile genome alignment system. PLoS Computational Biology 14:e1005944.

McKenna, A., M. Hanna, E. Banks, A. Sivachenko, K. Cibulskis, A. Kernytsky, K. Garimella et al. 2010. The Genome Analysis Toolkit: A MapReduce framework for analyzing next-generation DNA sequencing data. Genome Research 20:1297-1303.

Mölder, F., K. P. Jablonski, B. Letcher, M. B. Hall, P. C. van Dyken, C. H. Tomkins-Tinch, V. Sochat et al. 2025. Sustainable data analysis with Snakemake. F1000Research 10:33.

Msuthwana, P., Y. Zhou, Y. Song, Z. Feng, Y. Sui, J. Hu, and Y. Sun. 2022. Expression of Frizzled-4 Gene on Developing and Postnatal Anser anser and Anser cygnoides Feather Follicles. Indian Journal of Animal Research.

Nadeau, N. J., T. Burke, and N. I. Mundy. 2007. Evolution of an avian pigmentation gene correlates with a measure of sexual selection. Proceedings of the Royal Society B: Biological Sciences 274:1807-1813.

Nadeau, N. J., F. Minvielle, S. Ito, M. Inoue-Murayama, D. Gourichon, S. A. Follett, T. Burke et al. 2008. Characterization of Japanese quail yellow as a genomic deletion upstream of the avian homolog of the mammalian ASIP (agouti) gene. Genetics 178:777-786.

Pei, J., X. Pan, G. Wei, and Y. Hua. 2023. Research progress of glutathione peroxidase family (GPX) in redoxidation. Frontiers in Pharmacology 14:1147414.

Quinlan, A. R., and I. M. Hall. 2010. BEDTools: A flexible suite of utilities for comparing genomic features. Bioinformatics 26:841-842.

Raza, S. H. A., S. D. Pant, A. K. Wani, H. H. Mohamed, N. E. Khalifa, H. M. Almohaimeed, A. R. Alshanwani et al. 2022. Krüppel-like factors family regulation of adipogenic markers genes in bovine cattle adipogenesis. Molecular and Cellular Probes 65:101850.

Reed, K. M., K. M. Mendoza, J. E. Abrahante, S. G. Velleman, and G. M. Strasburg. 2021. Data Mining Identifies Differentially Expressed Circular RNAs in Skeletal Muscle of Thermally Challenged Turkey Poults. Frontiers in Physiology 12:732208.

Rius-Pérez, S., I. Torres-Cuevas, I. Millán, Á. L. Ortega, and S. Pérez. 2020. PGC-1α, Inflammation, and Oxidative Stress: An Integrative View in Metabolism. Oxidative Medicine and Cellular Longevity 2020:1-20.

Rong, H., X. He, L. Zhu, X. Zhu, L. Kang, L. Wang, Y. He et al. 2017. Association between regulator of telomere elongation helicase1 (RTEL1) gene and HAPE risk. Medicine 96:e8222.

Ruegg, K., E. C. Anderson, J. Boone, J. Pouls, and T. B. Smith. 2014. A role for migration-linked genes and genomic islands in divergence of a songbird. Molecular Ecology 23:4757-4769.

Saldana-Caboverde, A., and L. Kos. 2010. Roles of endothelin signaling in melanocyte development and melanoma. Pigment Cell & Melanoma Research 23:160-170.

Schweizer, T. M., and M. G. DeSaix. 2023. Cost-effective library preparation for whole genome sequencing with feather DNA. Conservation Genetics Resources 15:21-28.

Schwob, M. R., M. B. Hooten, and V. Narasimhan. 2024. Composite dyadic models for spatio-temporal data. Biometrics 80:ujae107.

Simonson, T. S., Y. Yang, C. D. Huff, H. Yun, G. Qin, D. J. Witherspoon, Z. Bai et al. 2010. Genetic Evidence for High-Altitude Adaptation in Tibet. Science 329:72-75.

Skoglund, P., and J. Höglund. 2010. Sequence polymorphism in candidate genes for differences in winter plumage between Scottish and Scandinavian Willow Grouse (Lagopus lagopus). PLoS One 5:e10334.

Smith, S., and N. Ellis. 2013. gradientForest: Random Forest functions for the Census of Marine Life synthesis project.

Trabucco, A., and R. J. Zomer. 2019. Global high-resolution soil-water balance, figshare.

Van der Auwera, G. A., M. O. Carneiro, C. Hartl, R. Poplin, G. Del Angel, A. Levy-Moonshine, T. Jordan et al. 2013. From FastQ data to high-confidence variant calls: The genome analysis toolkit best practices pipeline. Current Protocols in Bioinformatics 43:11-10.

Wan, Y., Z. Zhang, D. Lin, X. Wang, T. Huang, J. Su, J. Zhang et al. 2022. Characterization of CRH-binding protein (CRHBP) in chickens: Molecular cloning, tissue distribution and investigation of its role as a negative feedback regulator within the hypothalamus–pituitary–adrenal axis. Genes 13:1680.

Wang, D., L. Liu, Z. Qu, B. Zhang, X. Gao, W. Huang, M. Feng et al. 2022. Hypoxia-inducible factor 1α enhances RANKL-induced osteoclast differentiation by upregulating the MAPK pathway. Annals of Translational Medicine 10:1227-1227.

Wang, T., A. Hamann, D. Spittlehouse, and C. Carroll. 2016. Locally downscaled and spatially customizable climate data for historical and future periods for North America. PloS One 11:e0156720.

Wang, T., M. Zhou, J. Guo, Y.-Y. Guo, K. Ding, P. Wang, and Z.-P. Wang. 2021. Analysis of selection signatures on the Z chromosome of bidirectional selection broiler lines for the assessment of abdominal fat content. BMC Genomic Data 22:18.

Wang, W., Y. Li, Z. Li, N. Wang, F. Xiao, H. Gao, H. Guo et al. 2020. Polymorphisms of KLF3 gene coding region and identification of their functionality for abdominal fat in chickens. Veterinary Medicine and Science 7:792-799.

Yang, K., J. Zhang, Y. Zhao, Y. Shao, M. Zhai, H. Liu, and L. Zhang. 2023. Whole Genome Resequencing Revealed the Genetic Relationship and Selected Regions among Baicheng-You, Beijing-You, and European-Origin Broilers. Biology 12:1397.

Yi, X., Y. Liang, E. Huerta-Sanchez, X. Jin, Z. X. P. Cuo, J. E. Pool, X. Xu et al. 2010. Sequencing of 50 Human Exomes Reveals Adaptation to High Altitude. Science 329:75-78.

Yin, H., D. Li, Y. Wang, and Q. Zhu. 2019. Whole-genome resequencing analysis of Pengxian Yellow Chicken to identify genome-wide SNPs and signatures of selection. 3 Biotech 9:383.

Zhuang, Z.-X., S.-E. Chen, C.-F. Chen, E.-C. Lin, and S.-Y. Huang. 2020. Genomic regions and pathways associated with thermotolerance in layer-type strain Taiwan indigenous chickens. Journal of Thermal Biology 88:102486.
